# Stimulus information guides the emergence of behavior related signals in primary somatosensory cortex during learning

**DOI:** 10.1101/2022.12.04.518156

**Authors:** Mariangela Panniello, Colleen J Gillon, Roberto Maffulli, Marco Celotto, Stefano Panzeri, Blake A Richards, Michael M Kohl

**Affiliations:** Department of Physiology, Anatomy and Genetics, University of Oxford, Oxford, OX1 3PT, UK; School of Psychology and Neuroscience, University of Glasgow, Glasgow, G12 8QQ, UK; Optical Approaches to Brain Function Laboratory, Istituto Italiano di Tecnologia, Genova, Italy; Department of Biological Sciences, University of Toronto Scarborough, Toronto, Ontario, Canada; Department of Cell & Systems Biology, University of Toronto, Toronto, Ontario, Canada; Mila, Montréal, Québec, Canada; Neural Computation Laboratory, Center for Human Technologies, Istituto Italiano di Tecnologia, Genova, Italy; Department of Neural Information Processing, Center for Molecular Neurobiology (ZMNH), University Medical Center Hamburg-Eppendorf (UKE), Hamburg, Germany; Department of Pharmacy and Biotechnology, University of Bologna, Bologna, Italy; School of Computer Science, McGill University, Montréal, Québec, Canada; Department of Neurology & Neurosurgery, McGill University, Montréal, Québec, Canada; Learning in Machines and Brains Program, Canadian Institute for Advanced Research, Toronto, Ontario, Canada; Montreal Neurological Institute, Montréal, Québec, Canada

## Abstract

Cortical neurons in primary sensory cortex carry not only sensory but also behavior-related information. However, it remains unclear how these types of information emerge and are integrated with one another over learning and what the relative contribution of activity in individual cells versus neuronal populations is in this process. Current evidence supports two opposing views of learning-related changes: 1) sensory information increases in primary cortex or 2) sensory information remains stable in primary cortex but its readout efficiency in association cortices increases. Here, we investigate these questions in primary sensory cortex during learning of a sensory task. Over the course of weeks, we imaged neuronal activity at different depths within layers 2 and 3 of the mouse vibrissal primary somatosensory cortex (vS1) before, during, and after training on a whisker-based object-localization task. We leveraged information theoretical analysis to quantify stimulus and behavior-related information in vS1 and estimate how much neural activity encoding sensory information is used to inform perceptual choices as sensory learning progresses. We also quantified the extent to which these types of information are supported by an individual neuron or population code. We found that, while sensory information rises progressively from the start of training, choice information is only present in the final stages of learning and is increasingly supported by a population code. Moreover, we demonstrate that not only the increase in available information, but also a more efficient readout of such information in primary sensory cortex mediate sensory learning. Together, our results highlight the importance of primary cortical neurons in perceptual learning.

## INTRODUCTION

Neuronal activity in the vibrissal primary somatosensory cortex (vS1) of mice successfully trained on a sensory task reflects not only sensory stimuli but also various types of behavior-related information^1,2^, including information about behavioral choice^3–8^. Insight into learning-related changes in cortical neuronal activity is key to understanding how the brain enables flexible behavior. On an individual neuron level, a variety of learning-related changes have been observed in vS1, including sharpening of neuronal responses^3,9^ and changes in the magnitude of neuronal signals^7^. It has been theorized that such changes serve to increase the ability of neurons to discriminate between similar pieces of information, thereby improving behavioral performance on related tasks^10^. Yet, some studies report minimal changes in the response properties of individual vS1 neurons over the course of learning^5,11^ and instead find learning-related alterations at the population level, for example in the relative spike-timing^12^, in neuronal gain^3^, or in population activity correlations (for review, see ^13^). The field still lacks a comprehensive picture of how stimulus and behavior-related information emerge and are integrated with one another over time as learning takes place, and what the relative contribution of activity in individual cells *versus* neuronal populations is in this process. We hypothesized that task-learning is supported by gradual changes at the individual neuron and population levels, which result in both increased information about sensory stimuli, and a more efficient use of this information to guide behavior. We anticipate that this, in turn, contributes to generating novel, task-specific information, necessary for behavioral improvement. We tested this hypothesis by training mice on a head-fixed tactile object localization task^14,15^, using two-photon imaging to longitudinally record the activity of excitatory neurons at different depths within layers 2 and 3 (L2/3) of vS1 before, during and after training. We quantified, on a trial-by-trial basis and at different stages of learning, stimulus information (MI-RS), behavioral choice information (MI-RC), and intersection information (II). II quantifies the amount of sensory information carried in the neural response that is read out to inform behavioral choice and provides potential insight into how information encoding supports sensory-guided behavior. We revealed that stimulus information was already present at the beginning of training, while choice information only emerged over the course of learning. Furthermore, we found that the improvement in behavioral performance was not simply accompanied by increased stimulus information but that, across learning stages, this information was more efficiently read out to instruct behavior. Finally, while changes in sensory information content were mainly shaped by changes at the individual neuron level, an increase in information encoded at the neuronal population level was associated more with behavioral choice.

## RESULTS

### Multi-depth two-photon calcium imaging over the course of learning

We trained mice to learn a whisker-based object localization task^14,15^ while they were head-fixed but freely running on a cylindrical treadmill. Mice learned to report a Go or No-go position of a vertical metal pole presented against the left whiskers by licking for a water reward. Learning was classified into three stages, based on the percentage of correct licking responses: ≤55% (stage 1), >55 to ≤75% (stage 2), >75% (stage 3) (**Fig. 1a-c**). During each of the three learning stages, we recorded the responses of excitatory neurons at four depths in the supragranular portion of the vibrissal primary somatosensory cortex (vS1) which expressed the genetically encoded calcium indicator GCaMP6s^16^, using multi-depth two-photon calcium imaging^17^. Learning progress was monitored using lick events. Over the course of learning, the time until the first lick after stimulus offset decreased substantially during correct Hit, but not incorrect false alarm (FA) trials (Hits: from 1.25±0.06 to 0.52±0.02 seconds; Kolmogorov-Smirnov test (ks-test) p<0.001; FAs: from 1.13±0.95 to 1.31±0.08 seconds; ks-test p=0.011; mean; **Fig. 1d**). On average, mice took 10.4±0.9 days of training to reach learning stage 3 (**Fig. 1e**). The mean percentage of correct responses on the day of best performance was 82.7% across mice (sd: 4.06; mean d’: 2.31±0.47). In each animal, we recorded vS1 neuronal activity in the same four fields of views (FOVs) in layer 2 and 3 (L2/3) across training sessions (**Fig. 1f & g**). The overall number of neurons imaged over the course of learning stages (**Supp. Table 1**) as well as image quality (**Supp. Fig. 1**) remained stable.

**Figure 1:**
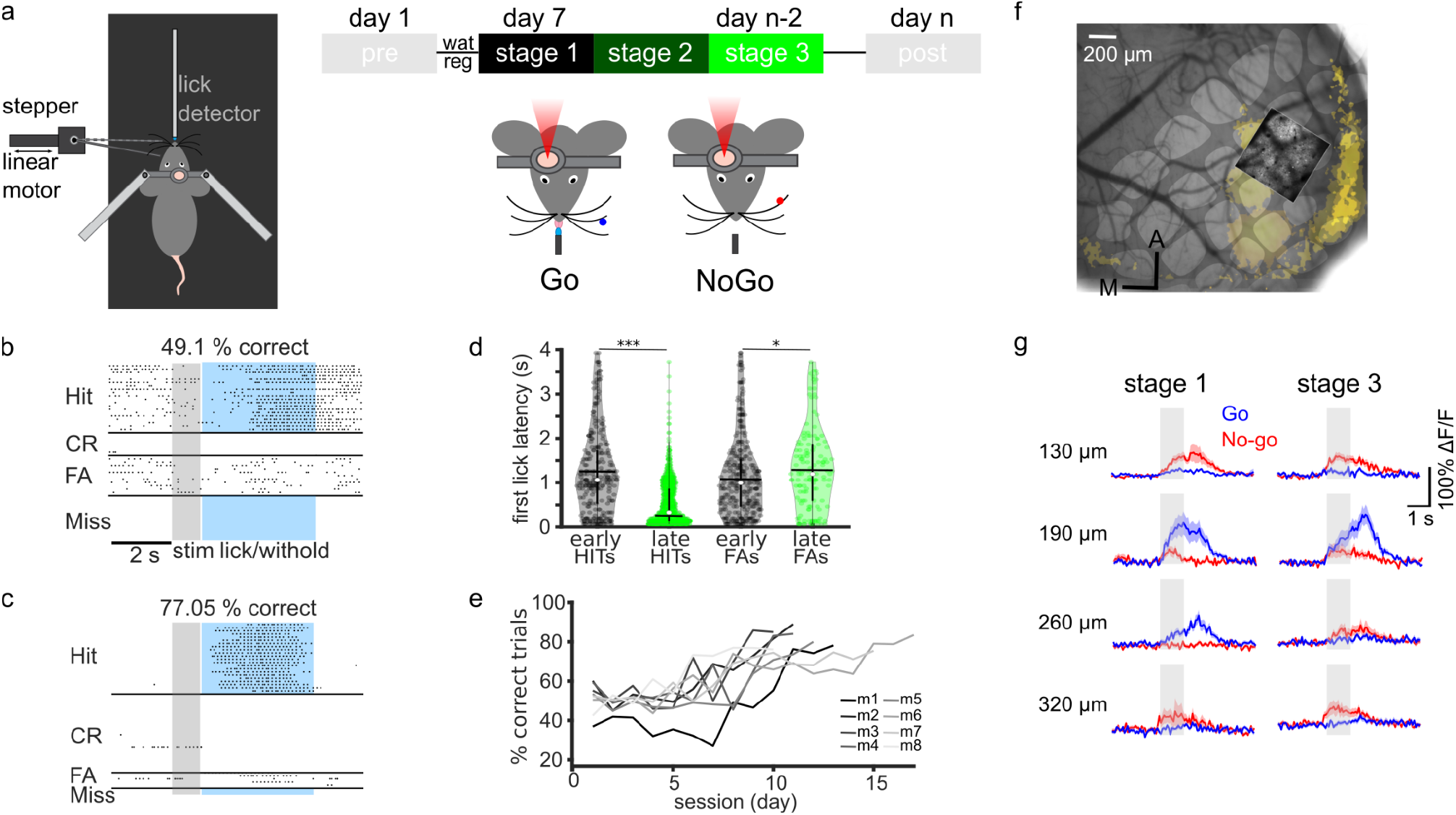
Learning of a tactile object localization task with concomitant vS1 calcium imaging. **a**. Experimental set up and protocol for imaging and sensory training. Mice were head-fixed, but free to run on a treadmill. On each trial, a metallic pole was moved toward the left whiskers to one of two positions (anterior: Go, and posterior: No-go). A spout was placed in front of the mice and used to deliver water when a lick was detected during a Go trial. Mice underwent water regulation before starting sensory training. Calcium transients in the right vS1 were recorded from the start of sensory training until the mouse achieved >70% correct responses for three consecutive days. Sensory learning was divided into 3 stages, based on their correct performance (stage 1, black: ≤55%; stage 2, dark green: >55%< to ≤75%; stage 3, light green: >75%). **b. & c**. Lick timings during each trial of an early training session (b., 49% correct), and a late training session (c., 77% correct). Trials are sorted according to trial outcomes. Trials where the mouse licked only during tactile stimulation were excluded from the plot and from the analysis. The gray shaded area indicates the time during which the pole was in contact with the whiskers. The blue shaded area indicates the 4-seconds-long licking window. Correct responses included licking on Go trials (Hit), and withholding licks on No-go trials (correct rejection, CR). Incorrect responses included withholding licks on Go trials (Miss), and licking on No-go trials (false alarm, FA). **d**. First lick latencies during Hit trials and FA trials for early training sessions (performance <55% correct; black data points, n=286 licks for Hits, 303 licks for FAs, across 8 mice) and late training sessions (performance >75% correct; green data points, n=949 licks for Hits, 150 licks for FAs, across 8 mice). Latency is calculated from stimulus offset. White dots indicate the median of the distribution, horizontal black bars indicate the mean value, vertical black bars indicate the 25th and 75th percentiles. **e**. Fraction of correct responses ((Hits + CR) / total trials) in all 8 mice, across up to 17 days of training. **f**. A representative IOSI image showing the location of the barrels corresponding to the whiskers stimulated during the procedure (yellow shading) in one mouse. A projection of the two-photon imaging FOV acquired throughout learning is overlaid on the IOSI image. Scale bar indicates 200 μm. g. Mean ΔF/F0 across Go trials (red) and No-go trials (blue) for one example neuron at each of the four cortical depths imaged. In this example, the same neuron was imaged during stages 1 and 3 of learning.

### Individual neurons in vS1 gain both stimulus and choice information over the course of learning

Neuronal responses to whisker touch were variable within individual FOVs, both in terms of stimulus preference (Go *vs*. No-go positions) and timing (early *vs*. late responses) at all learning stages (**Fig. 2a**). To quantify how much information about stimulus position was carried in the activity of each imaged neuron at each time point during a trial, we calculated instantaneous Mutual Information (MI) between a neuron’s calcium response and stimulus identity across trials (MI-RS; **Fig. 2b**). We repeated the process for each depth and learning stage. When averaging the frame-by-frame MI-RS values obtained for each imaged neuron, we found that the overall MI-RS increased across the learning stages at all imaging depths (ks-test p: all <0.001; **Fig. 2c**). MI-RS was already present during learning stage 1, as might be expected for a primary sensory region (p<0.001 when compared to the null distribution, for all depths). The percentage of neurons carrying significant MI-RS was very similar across cortical layers (−130 μm: 24.7%, -190 μm: 25.5%, -260 μm: 32.6%, -320 μm: 25.7%; **Fig. 2d**). MI-RS of individual neurons was higher for superficial than deep layers, but there was 4-fold increase in MI-RS at -320 μm between stage 1 and stage 3 of learning (MI-RS stage 1/MI-RS stage 3 at -130 μm: 2.46, -190 μm: 3.08, -260 μm: 2.37, -320 μm: 3.88), and a doubling of the number of neurons carrying significant MI-RS (−130 μm: 50.92%, -190 μm: 59.34%, -260 μm: 62.22%, -320 μm: 53.23%). Interestingly, MI-RS values showed a steep increase right after stimulus onset (defined as the moment when the metal pole reached its final position against the whiskers) but kept increasing after stimulus offset (defined as the moment when the metal pole started moving away from the whiskers), suggesting that stimulus information builds and persists in vS1 even when the stimulus is not present anymore.

**Figure 2:**
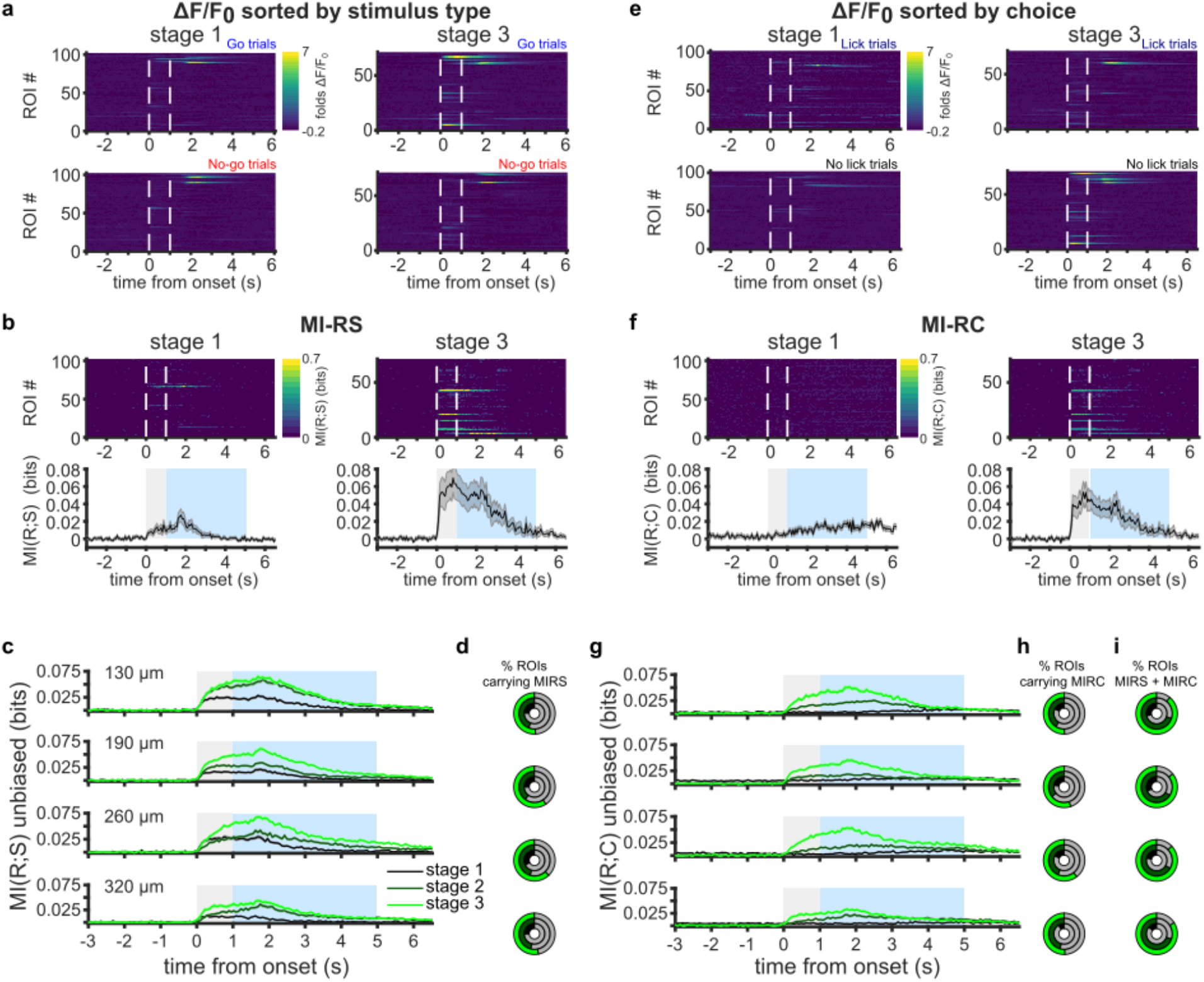
Stimulus and choice information increase over learning. **a**. (Left) Frame-by-frame mean ΔF/F_0_ across Go trials (top) and No-go trials (bottom) for all neurons (ROI #) in one example FOV, -190 μm below the cortical surface. Imaging was performed during learning stage 1. White lines indicate stimulus onset and offset. (Right) ΔF/F_0_ activity from the same FOV, but when the mouse was in learning stage 3. **b**. Left: Frame-by-frame mutual information between stimulus and response (MI-RS) in the same neurons represented in a., during stage 1. Right: MI-RS from the same FOV, when the mouse was in stage 3. Bottom left and right show the average MI-RS across all neurons in the field. The gray shaded area indicates stimulus duration. The blue shaded area indicates the licking window. **c**. Mean frame-by-frame MI-RS across all neurons (n=8 mice) at each cortical depth and for each learning stage. MI-RS was first averaged framewise across all neurons in the same FOV, and then averaged across all FOVs imaged at the same cortical depth and during the same learning stage (stage 1: black, stage 2: dark green, stage 3: light green). **d**. Fraction of neurons showing significant MI-RS at each cortical depth and at each learning stage. Learning stage is indicated by the color inside each concentric circle. Full circles correspond to 100% of imaged neurons. The gray shaded area in the circles indicates the fraction of neurons with non-significant MI-RS (p>=0.05). The remaining portion indicates significant neurons. **e**. ΔF/F_0_ activity for the same neurons as shown in a. Here, responses were separated into Lick (top) *vs*. No-lick trials (bottom). **f**. Same as in b, but for mutual information between response and mouse choice (*i*.*e*., Lick *vs*. No-lick, MI-RC). **g**. Same as in c. and d., but for MI-RC. **i**. Fraction of neurons carrying both significant MI-RS and significant MI-RC, calculated over the total number of neurons carrying significant MI-RS.

We next asked whether individual neurons in vS1 also represent the behavioral choice to lick or withhold licking, and whether this representation changes with learning. We therefore assessed MI between neural responses and choice (MI-RC), as above (**Fig. 2e & f**). Similar to MI-RS, we found that MI-RC increased across learning stages. However, whereas MI-RS was already present in learning stage 1, the mean MI-RC across imaged neurons was near zero early during training, but progressively increased through the following learning stages (MI-RC stage 3/stage 1 at -130 μm: 10.85, -190 μm: 4.17, -260 μm: 8.84, -320 μm: 5.26; **Fig. 2g**). This trend is reflected in the lower fraction of neurons carrying significant MI-RC, compared to MI-RS, in stage 1 (−130 μm: 18.1%, - 190 μm: 22.86%, -260 μm: 18.29%, -320 μm: 9.6%), increasing to more than half of the imaged neurons at stage 3 (−130 μm: 50.39%, -190 μm: 55.31%, -260 μm: 60.58%, -320 μm: 50.74%; **Fig. 2h**).

Moreover, by stage 3, the majority of neurons carrying significant MI-RS also showed significant MI-RC (**Fig. 2i**), hinting at a computation taking place during learning, where primary cortical neurons encoding sensory stimulus information are recruited to inform behavioral choice as well, and contribute to task performance.

Over the course of learning, information about the tactile stimulus and behavioral choice increased as mice improved their behavioral performance. Two notable cortical layer differences could be observed: First, stimulus information stops increasing during stage 2 in superficial L2 (−130 μm) and deep L3 (−320 μm). Second, the increase in stimulus information is strongest in deep L3 while the increase in choice information is strongest in superficial L2. Overall, these results show that, at the start of sensory training, stimulus information is already present, particularly in superficial layer 2 neurons, while choice information is absent.

### Learning-related increase in choice information is supported by population coding

Neurons in the same brain region vary in how strongly they encode sensory stimulus information, and neuronal activity in vS1 tends to be particularly sparse^18–20^. Thus, we next sought to evaluate how the learning-related changes in stimulus and choice information across neurons in vS1 reflect the contribution of individual neurons to the neuronal population encoding as a whole. Calculating MI on the activity of increasing numbers of individual neurons is subject to a systematic bias due to the limited number of experimental trials available^21^. Therefore, following established practices, we estimated the MI for groups of neurons using the MI computed on the confusion matrices obtained by training linear regression models to decode stimulus (decMI-RS) or choice (decMI-RC) from neural activity. decMI-RS and decMI-RC computed for groups of neurons offer a lower-bound to the amount of stimulus and choice information encoded by the neural population^22^. When computed for individual neurons, decMI-RS and decMI-RC correlated well with the previously estimated MI-RS and MI-RC, indicating that the measure was reliable (**Supp. Fig. 2**).

We first sought to use decMI to evaluate the contribution of each neuron to population-level encoding of task-relevant information. We classified each neuron as “discriminative” if it carried sufficient decMI on its own to enable above-chance decoding of the trial type (*i*.*e*., above the 95^th^ percentile of the null distribution) and “non-discriminative” otherwise^15^. We then evaluated whether the increase in stimulus and choice information over the course of learning reflected a population level change or the emergence of a sparse set of highly-informative discriminative neurons. Over the course of learning, median decMI-RS did not change but the percentage of discriminative neurons increased (stage 1: 16.0±1.2%, stage 2: 19.5±1.1%, stage 3: 24.4±2.2%; mean±sem across depths and FOVs) and the distribution of decMI-RS values changed, reflecting a change in the 95^th^ percentile (stage 1: 0.31, stage 2: 0.39, stage 3: 0.47; **Fig. 3a**). Similarly, for decMI-RC the percentage of discriminative neurons increased (stage 1: 7.4±1.0%, stage 2: 11.9±0.8%, stage 3: 19.6±2.0%) and the distribution of decMI-RS values changed (95^th^ percentile in stage 1: 0.21, stage 2: 0.25, stage 3: 0.32; **Fig. 3b**). Together, these results indicate that the increase in stimulus and choice information observed in L2-3 of vS1 reflects both an increase in the number of discriminative neurons, as well as an increase in the information about stimulus and choice carried by the most informative neurons.

**Figure 3:**
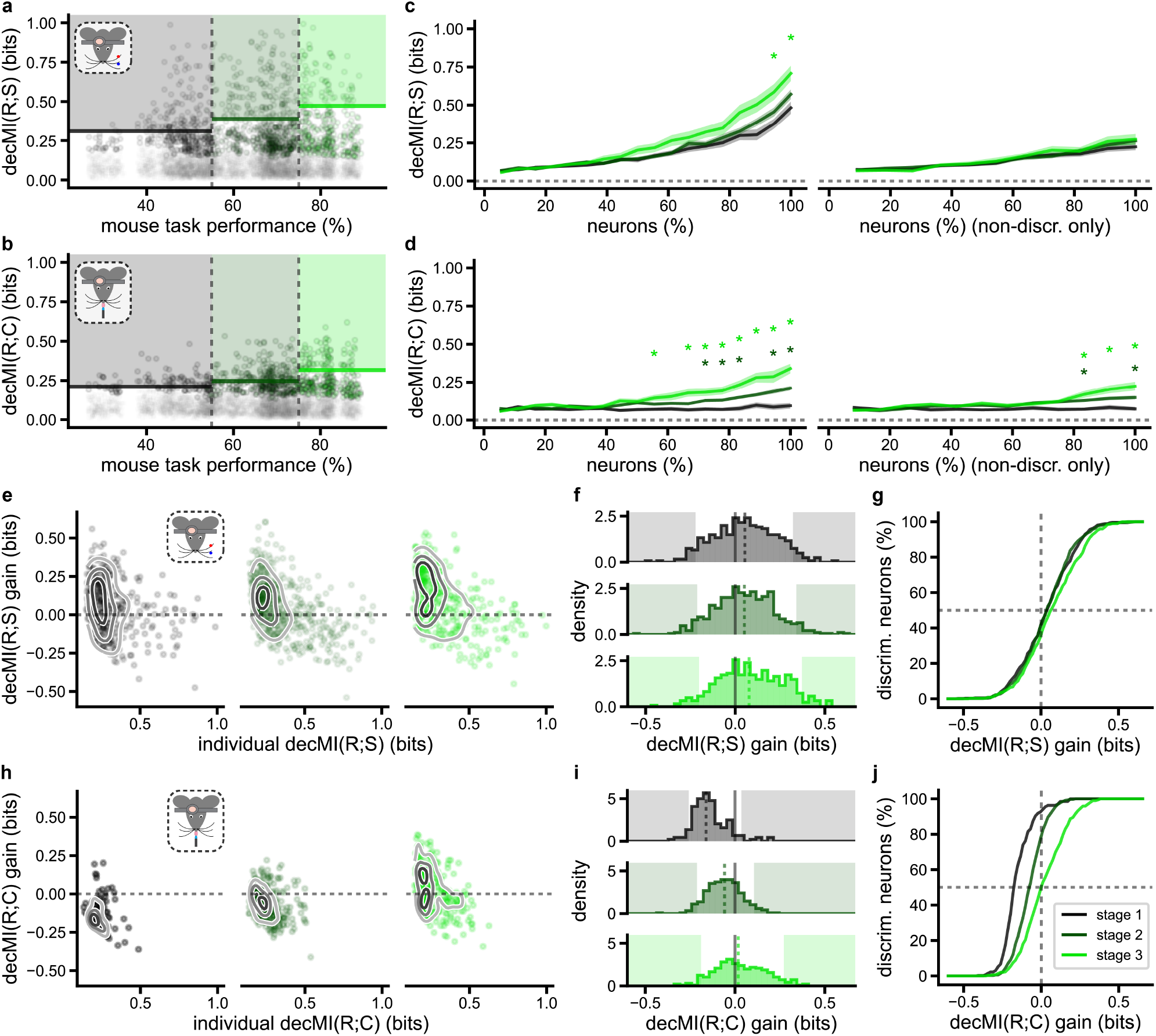
Population codes contribute more strongly to choice than stimulus information over learning. **a**. decMI-RS (mutual information calculated on stimulus decoding confusion matrices) plotted for individual neurons, against mouse task performance (plotted with some jitter in x). The learning stages are delineated by color and vertical black dashed lines. Only discriminative neurons (decMI-RS p<0.05, compared to the null distribution) are plotted in the learning stage color (stage 1: black, stage 2: dark green, stage 3: light green), with non-discriminative neurons plotted in gray (decMI-RS p>=0.05). Colored shading indicates values above the 95th percentile (indicated by bold line). **b**. Same as a., but for decMI-RC (MI calculated on behavioral choice decoding confusion matrices). **c**. Mean across FOVs of population decMI-RS as neurons are added to the pool fed to the decoder, in order of lowest to highest individual decMI-RS. The full pool includes either all neurons (left) or only non-discriminative neurons (right) (stage 1: black, stage 2: dark green, stage 3: light green). Asterisks indicate a significant difference between stage 1 and either stage 2 (dark green) or stage 3 (light green) (t-test p<0.05 corrected, as elsewhere, for multiple comparisons). The horizontal dashed line marks zero decMI-RS. **d**. Same as c., but for dec MI-RC. **e**. Discriminative neuron decMI-RS plotted against the gain in decMI-RS with respect to individual decMI-RS, when decoders also received non-discriminative neuron responses as input. Contour lines qualitatively show data density levels. Stages 1, 2 and 3 are plotted left to right. **f**. Histograms of decMI-RS gain (y axis data from e.), with stages 1, 2 and 3 plotted top to bottom. The black lines mark 0 gain, whereas median gain is indicated by a dashed line, and shaded areas show values below the 5^th^ or above the 95^th^ percentile of the distribution. **g**. Data from f., represented as cumulative sums. The vertical dashed line marks zero gain, whereas the horizontal dashed line marks the median of each distribution. As shown by the legend in j., data for each learning stage is plotted in the stage’s color (listed in a.). **h-j**. Same as e-g., but for decMI-RC (choice).

Previous work has shown that neurons which on their own do not enable above-chance decoding of task-relevant variables, can still contribute to population encoding and improve the decoding performance of neurons that carry high information content, when put together^23^. This points to a role for non-discriminative neurons in supporting robust population codes for task-relevant information. We wanted to determine the relative importance of these non-discriminative neurons for stimulus and choice information. We therefore asked how information about stimulus and choice increased as we added neurons, from least to most informative, to the pool used for calculating decMI. For each session, we ran decoders sequentially as we added neurons with progressively increasing decMI, drawn from either the full population (**Fig. 3c & d**, left) or only from the non-discriminative neuron population (**Fig. 3c & d**, right). We then compared decMI-RS in stages 2 and 3 to stage 1 values, as neurons were added, to identify differences between stages (t-test p<0.05, Bonferroni corrected for all neuron % × stage comparisons). When it came to decMI-RS, we found that as neurons were added to the pool, decMI-RS tended to increase for all stages. Only once 90% or more of the entire population of recorded neurons was included, decMI-RS was substantially higher for stage 3 FOVs compared to stage 1 FOVs, and no differences were found for stage 2 *vs*. 1 FOVs (**Fig. 3c**, left). No differences emerged between the stages when only non-discriminative neurons were included (**Fig. 3c**, right). In contrast, during stage 1, decMI-RC remained low regardless of the number of neurons added to the decoder (**Fig. 3d**). In stages 2 and 3, however, decMI-RC increased beyond stage 1 levels once 70% of all neurons were included in stage 2, and with as few as 55% of all neurons in stage 3. In contrast to decMI-RS, even adding only non-discriminative neurons significantly increased decMI-RC in stages 2 and 3 compared to stage 1 (**Fig. 3d**). Overall, these results suggest that, as mice learn to perform the task, choice information is increasingly supported by a distributed population code. In contrast, stimulus information shows a consistent reliance on a distributed population code across learning, that is already present at the start of training.

Finally, we wanted to quantify how much stimulus or choice information individual discriminative neurons gain from the activity of a population of non-discriminative neurons. We calculated the decMI of each discriminative neuron on its own and then measured the gain in information when the decoder also received as input the neuronal activity from the session’s non-discriminative neurons. Including the non-discriminative population greatly increases the dimensionality of the input to the decoders which, if the added input data is not informative, can impair a decoder’s performance. For stimulus decoding, this was the case for approximately half of all discriminative neurons which showed a negative gain when paired with the non-discriminative population, as shown by median gains near 0 (0.05 in stage 1, 0.05 in stage 2, 0.08 in stage 3; **Fig. 3e-g**). The discriminative neurons that showed positive gains were generally a subset of the ones that had the lowest individual decMI-RS (<0.5; **Fig. 3e**). The overall distribution of the gains in stimulus decoding showed negligible change across learning (**Fig. 3e-g**). In contrast, non-discriminative neurons had a much stronger effect on choice decoding by discriminative neurons. The median gain increased steadily with learning (stage 1: -0.16, stage 2: -0.06, stage 3: 0.02; **Fig. 3h-j**). The overall distribution of the gains in choice decoding showed a strong overall rightward shift (ks-test p<0.001 for all pairs of stages), with the 95^th^ percentile increasing substantially across stages (stage 1: 0.04, stage 2: 0.11, stage 3: 0.27). As for stimulus decoding, the discriminative neurons that gained the most from being paired with the non-discriminative population, across all stages, were generally those with lower individual decMI-RC (**Fig. 3h**). Together, these results suggest that, in vS1, over learning, neurons that encode task-relevant variables well on their own become progressively more supported in population encoding by neurons that would not be significantly discriminative on their own, particularly for choice encoding.

### Stimulus information increasingly guides behavioral choice throughout learning

The increase in perceptual abilities when learning a sensory-guided task may be due, as traditionally hypothesized, to an increase of the sensory information encoded in early sensory cortices^5,24–26^. Alternatively, it may be the consequence of an improved readout of such information^27^. To gain insights into how sensory information encoded in vS1 is used to generate accurate behavior across stages of learning, we used intersection information (II)^28,29^, an information-theoretic quantification of how much sensory information in neural activity is read out to inform behavioral choices (**Fig. 4a**). By definition, II is non-negative, on a scale of bits, and is bounded by both MI-RS and MI-RC. First, we calculated the frame-by-frame II carried by each imaged neuron, across trials, at each depth and learning stage. II was, as expected, absent before stimulus onset, at all learning stages, because there was no stimulus information during this time window. After stimulus-onset, II was weak during learning stage 1, but increased in stages 2 and 3 (**Fig. 4b**). The percentage of neurons carrying significant II was low in all recorded layers in stage 1 but increased to more than half of the imaged neurons in stage 3, irrespective of cortical depth (−130 μm: 52.0%, -190 μm: 57.1%, -260 μm: 58.7%, -320 μm: 50.9%; **Fig. 4c**). The emergence of II may be the result of two processes: (1) the increase in sensory information (MI-RS) available in neural activity over learning (**Fig. 2**) or (2) an increase in the efficiency by which sensory information stored in vS1 is read out downstream to inform behavior. To determine the relative contribution of either process, we calculated the ratio of II/MI-RS for each neuron carrying significant II, at each depth and learning stage. This ratio quantifies the proportion of sensory information available in neural activity that is actually read out to inform sensory behavior. We found that II/MI-RS increased over learning in all recorded cortical depths, peaking at ratios >0.75 in stage 3 (**Fig. 4d & e**). Importantly, all these findings also held when controlling for increased correlation between stimulus and choice across learning by trial stratification (**Supp. Fig. 4**.) In summary, during learning stage 1, some stimulus information is present but very little of it is read out. The increase in object-localization performance across learning is accompanied not only by an increase of sensory information available in the neural activity of vS1, but also by an increase in the efficiency by which this sensory information is read out to inform behavioral choices. By learning stage 3, more than 75% of the MI-RS could be used to guide the animal’s behavioral choice. These results were confirmed when using a simple decoder analysis^30,31^ (**Supp. Fig. 3**). This suggests that increased object-localization accuracy with learning is mediated by both the traditionally posited increase of sensory information in sensory areas and, more surprisingly, by an increased efficiency of the downstream readout of this information.

**Figure 4:**
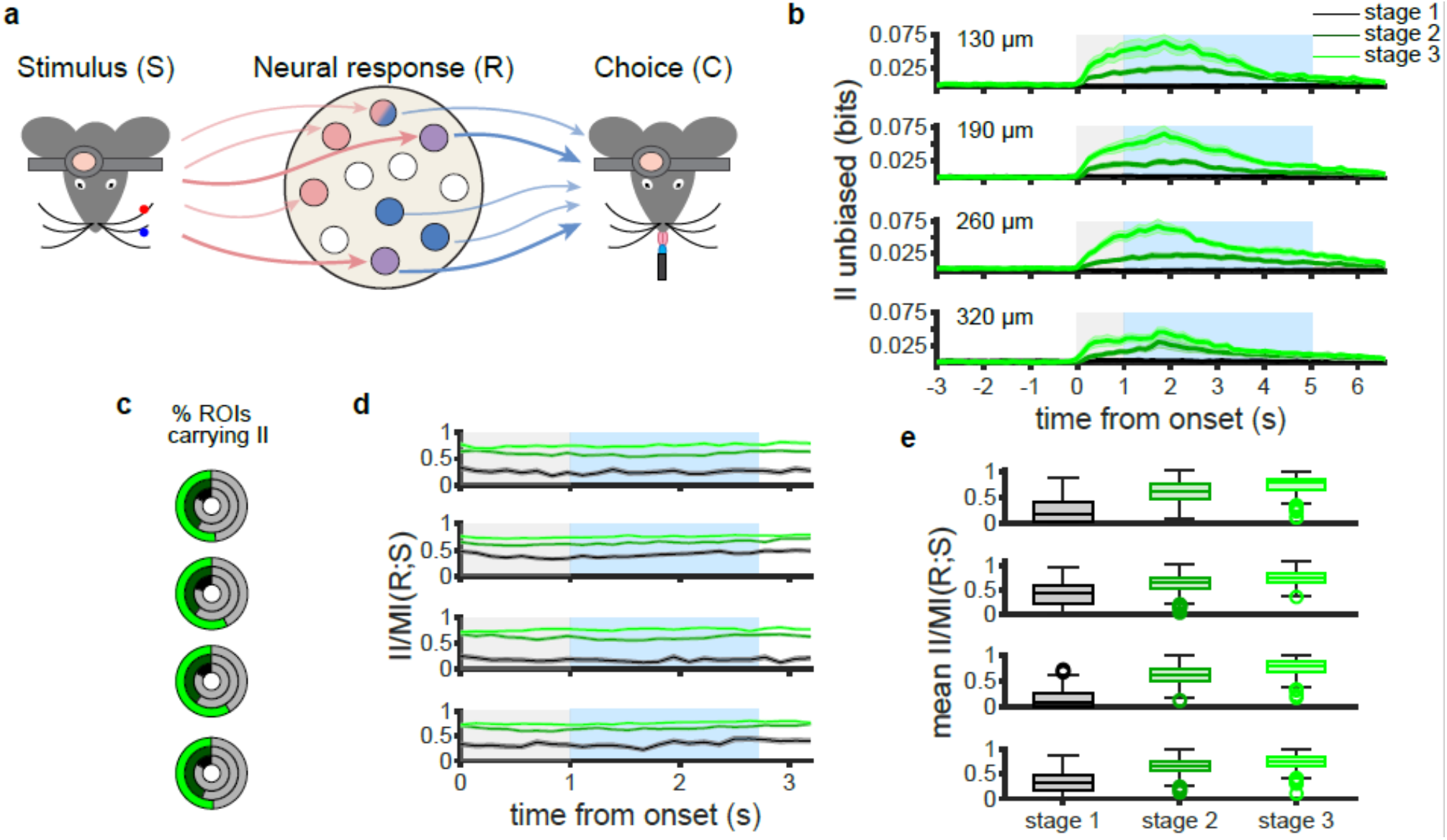
Contribution of stimulus information to behavioral choice. **a**. Schematic representation of the information theoretic framework showing the 2 stages of information processing. Stimulus encoding represents the mapping of the tactile stimuli onto the responses of neurons in L2-3 of vS1. Information readout is represented by the mapping of neuronal activity onto the mouse choice to lick or withhold licking in the presence of the tactile stimuli. Neurons, represented as circles, carry only MI-RS (pink) only MI-RC (blue), neither (white) or both. The latter neurons are represented as half blue, half pink, if they carry both MI-RS and MI-RC, but not intersection information (II). However, they are represented as purple if they carry II, with thick arrows coming in from the stimulus and going out to the choice, as these neurons carry stimulus information that directly informs mouse choice. **b**. Mean frame-by-frame II across all neurons (8 mice) at each cortical depth and for each learning stage. Gray shading indicates stimulus duration. Blue shading indicates the duration of the licking window. II was first averaged framewise across all neurons in the same FOV, and then averaged across all FOVs imaged at the same cortical depth and during the same learning stage. Learning stages are color coded (stage 1: black, stage 2: dark green, stage 3: light green). **c**. Fraction of neurons carrying significant II at each cortical depth and at each learning stage. Learning stage is indicated by the color inside each concentric circle. Full circles correspond to 100% of imaged neurons. The gray area in the circles indicates the fraction of neurons with non-significant II (p>=0.05). The remaining portion indicates neurons with significant II. **d**. Frame-by-frame II/MIRS for all the neurons that showed significant II, at each cortical depth and during each learning stage. Color code is the same as in b and c. **e**. Mean II/MIRS across frames for all neurons with significant II for each learning stage and cortical depth. Empty dots are outlier neurons. Horizontal lines indicate the median, and error bars indicate the lower and upper quartiles.

### Correct readout of stimulus information in neural activity developed during learning is altered by rule switch

We next sought to investigate how changing the relationship between stimulus and reward after learning affects neuronal encoding of stimulus and choice information. To do this, we recorded a “switch day” in three mice that had achieved >70% correct trials over three consecutive days. On this switch day, the task rule was reversed, and mice received a water reward if they licked when the stimulus was in the No-go position, while no water was delivered when mice licked in the Go position (**Fig. 5a**). Mouse performance during the switch day dropped to 37.9±3.5% (mean±sd).

**Figure 5:**
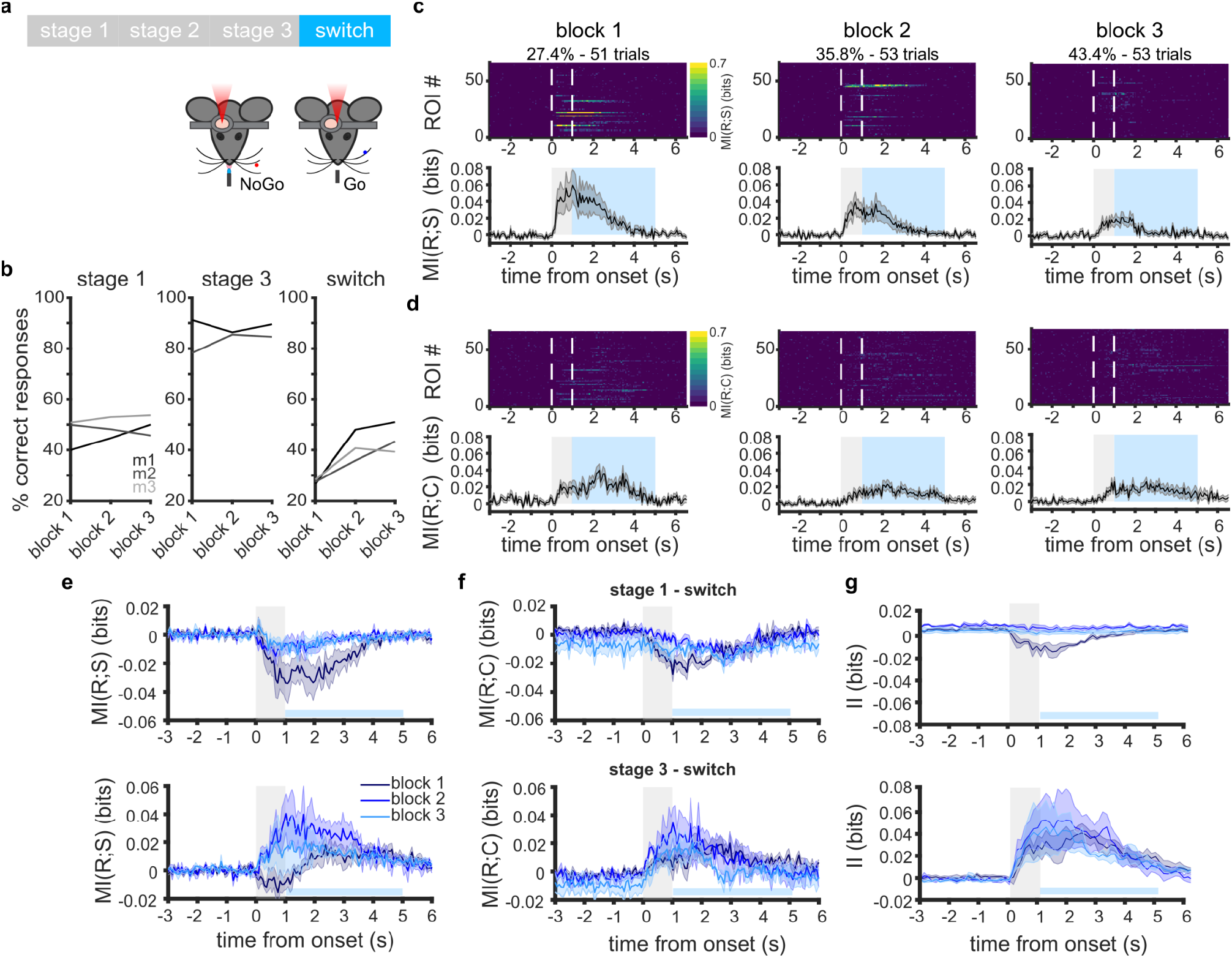
Choice information decreases before stimulus information when stimulus-reward association is altered. **a**. Experimental protocol. After reaching behavioral threshold (performance over 70% correct trials for three consecutive days) vS1 activity was imaged while 3 mice were tested on a “switch day”. On that day, the Go and No-go contingencies were switched. **b**. Behavioral performance of the three individual mice used for the switch sessions (mouse 1: black, mouse 2: dark gray, mouse 3: light gray), separately for each of the 3 blocks, during stage 1 (left), stage 3 (center) and switch (right) sessions. **c**. MI-RS for all neurons in one example FOV, during a switch session. MI-RS is shown frame-by-frame, for each putative neuron (top), and as the mean across neurons (bottom). Above the plots, the behavioral performance of the mouse is reported as the % of correct responses, for each of the three imaging blocks of the switch session. White lines in the top row indicate stimulus onset and offset. The gray and blue shaded areas in the bottom row indicate stimulus duration and licking window duration, respectively. **d**. Same as in c, but for MI-RC. **e-g**. Top: for each of the 3 blocks, the MI-RS, MI-RC and II computed during the switch session were subtracted respectively from the MI-RS, MI-RC and II obtained in the same FOV during learning stage 1. Data show mean across FOVs in three mice. Bottom: same, but in this case MI-RS, MI-RC and II computed during the switch session were subtracted from the values obtained in the same FOV during stage 3.

Mean MI-RS and MI-RC also dropped but stayed above the levels seen during stage 1 (**Supp. Fig. 4)**. We wondered about the time course of this drop in information relative to the drastic changes in behavioral performance during the switch. We therefore considered behavioral performance, MI-RS and MI-RC separately for each of the three daily training blocks. During learning stages 1 and 3, behavioral performance was stable across the three daily training blocks. In contrast, performance collapsed to very low levels in the first block of the switch, recovering to chance levels by the last, third block (**Fig. 5b**). Concurrently, MI-RS remained relatively high during block 1, and decreased gradually over the course of the switch session, whereas MI-RC was low throughout (**Fig. 5c**). We next sought to assess the difference in stimulus and choice information between the individual blocks of the switch session and learning stages 1 and 3. We therefore subtracted, for each FOV, the frame-by-frame MI values of the switch session from the MI values of learning stages 1 and 3. MI-RS_switch_ was overall higher than MI-RS_stage1_ throughout the switch session, and especially during block 1 (**Fig. 5e**, top), although the behavioral performance of each of the three mice was lower in this block than during stage 1 (**Fig. 5b**, left and right). MI-RS_switch_ was closer to MI-RS_stage1_ during blocks 2 and 3. When comparing MI-RS_switch_ to MI-RS_stage3_, MI-RS_switch_ was still slightly higher during block 1 (**Fig. 5e**, top), despite the lower behavioral performance (**Fig. 5b**, center and right). In contrast, MI-RS_switch_, which remained stable in blocks 2 and 3, was lower during these blocks than it had been in stage 3. These results show that learning the association between a stimulus and a reward strengthens the former’s representation in layer 2-3 vS1 (MI-RS_switch_ was initially higher than MI-RS_stage1_). Moreover, this representation remains stable only briefly (in our paradigm, about 50 trials) after the association with the reward has been altered (MI-RS_switch_ was initially slightly higher than MI-RS_stage3_), in a way that is independent from behavioral performance (*i*.*e*., independent from the animal being able to fully recognize the change in association). On the other hand, information about choice followed a different time course, showing a faster drop during the switch session. Whereas MI-RC_switch_ was slightly higher than MI-RC_stage1_ during all 3 blocks (**Fig. 5f**, top), it dropped below MI-RC_stage3_ as early as in block 1 (**Fig. 5f**, bottom). When measuring II_switch_, we found that it was still present in vS1 at the start of the switch session. This indicates that during block 1 of the switch session stimulus information was still read out to guide behavior, but the readout was incorrect (as performance dropped below chance level). During block 2 post switch, II_switch_ and II_stage1_ were both near zero, indicating that, at that point stimulus information was no longer used to inform choice. As expected, II_switch_ was lower than II_stage3_ during all 3 blocks of the switch session (**Fig. 5g**). These results show that information about stimulus and choice decrease but remain present throughout the switch session. However, their relationship changes after the switch. Stimulus information did not inform behavioral choice and became rapidly decoupled from it after the association stimulus-reward was altered, as intersection information disappeared. Together, these results further underline that the efficient and correct readout of stimulus information, built during sensory training, is key to the learning of object-localization tasks.

### Task-learning produces generalized and persistent increase in information

We have so far described the changes in information present in cortical circuits that occur when sensory stimuli are associated with a reward. To conclude, we wanted to know whether these learning-related changes in information generalize to stimuli not used in the task and persist without reward.

In seven of the eight mice trained on the object localization task, we imaged activity in L2-3 vS1 neurons during two further sessions (“pre-training” and “post-training”) in which sensory stimuli were presented outside of the context of the Go/No-go task, *i*.*e*., without the spout to lick or the associated water reward. The pre-training session was performed before water regulation and task training started, while the post-training session took place two days after the end of the task training. The stimulus was now presented in six different positions of which position 3 and 6 corresponded to the Go and No-go cue positions used during task training (**Fig. 6a**). To find out how much information about stimuli 3 and 6 was present in neurons of vS1 before and after training, we calculated the frame-by-frame MI-RS carried by each imaged neuron, across trials, at each depth and learning stage. Average MI-RS across neurons was low at all cortical depths when the mice experienced the whisker stimulation for the first time (*i*.*e*., during the pre-training session). Stimulus information more than doubled after training and the fraction of neurons carrying significant MI-RS increased in all cortical layers (**Fig. 6b & c**). The fraction of neurons carrying significant MI-RS during the pre-training session was consistently lower than the fraction of neurons carrying significant MI-RS during stage 1 of training (**Fig. 2d**), suggesting that more neurons are recruited to encode stimulus information as soon as the stimulus-reward association is introduced.

**Figure 6:**
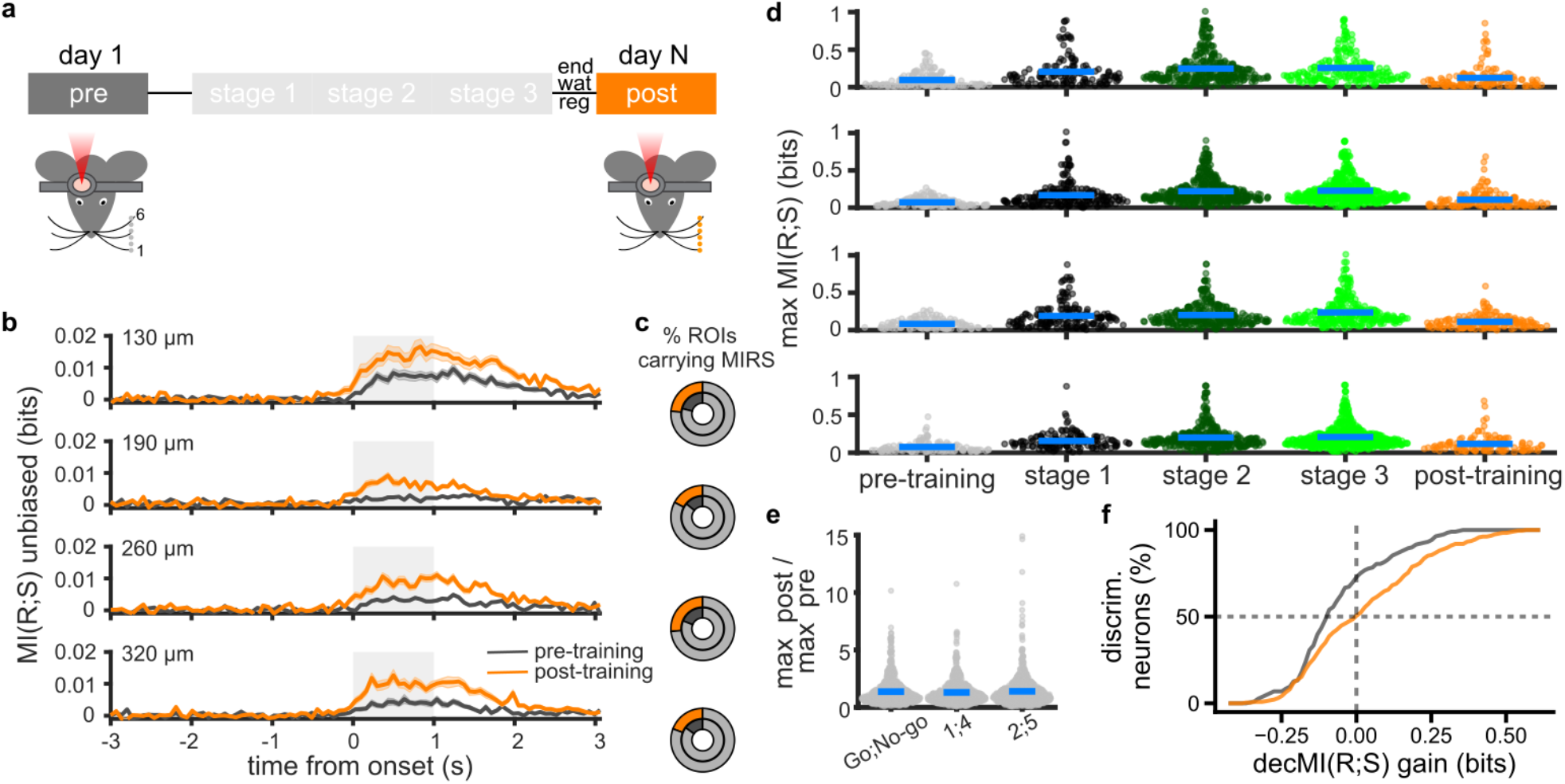
Sensory information persists in vS1 after learning. **a**. Schematic representation of the experimental protocol (n=7 mice). ΔF/F_0_ was measured for putative neurons in vS1 while mice were presented with a metallic pole in six different positions along their anterior-posterior axis, both before (pre-training session, gray) and after training on the pole localization task (post-training, orange). **b**. Mean frame-by-frame MI-RS across all neurons imaged during the pre-training (gray) and post-training (orange) sessions, for each frame and for each cortical depth (−130 μm: 659 (pre) and 527 (post) neurons, -190 μm: 879 (pre) and 765 (post) neurons, -260 μm: 554 (pre) and 557 (post) neurons, -320 μm: 747 (pre) and 567 (post) neurons). MI-RS was calculated on responses to stimulus positions 3 and 6 only (*i*.*e*., the pole positions used for sensory training). It was first averaged framewise across all neurons in the same FOV, and then across all FOVs imaged at the same cortical depth and during the same session. The gray shaded areas indicate stimulus duration. **c**. Fraction of neurons carrying significant MI-RS at each cortical depth (p<0.05). Full circles reflect 100% of imaged neurons. The light-gray area in each circle indicates the fraction of neurons with non-significant MI-RS. The dark-gray and orange portions indicate the fraction of neurons with significant MI-RS during the pre-training and post-training sessions, respectively. **d**. Distribution of the maximum MI-RS value (across frames) for each putative neuron with significant MI-RS. Data are shown for each depth, and for each passive and active imaging session: pre-training (gray), stage 1 training (black), stage 2 training (dark green), stage 3 training (light green), post-training (orange). Light blue bars indicate the mean value for each distribution. **e**. Ratio between maximum MI-RS value during post-training and the maximum MI-RS value during pre-training, calculated on each putative neuron that was tracked across the two imaging sessions (n=849 neurons in seven mice). MI-RS and ratios were calculated separately for stimuli 3 and 6 (also used during training, left), for stimuli 1 and 4 (center), and for stimuli 2 and 5 (right). Data were pooled across cortical depths. Blue lines indicate the mean for each distribution. **f**. Cumulative sums of gain in decMI-RS observed for each discriminative neuron when decoders also received non-discriminative neuron responses as input. The vertical dashed line marks zero gain, whereas the horizontal dashed line marks the median of each distribution. As in panels b. and c., the data for pre and post training are plotted in gray and orange, respectively.

We then asked whether sensory information improved specifically for the stimulus positions used in the object localization task, or whether the MI-RS increase reflected a general increase in object location information in vS1. We computed MI-RS on pairs of pole positions (1 and 4, 2 and 5) separated by the same distance as the Go / No-go positions 3 and 6. Before training, the percentage of significant MI-RS neurons was comparable across pairs of stimuli (mean across layers for 3 and 6: 17.3%; 1 and 4: 14.8%; 2 and 5: 16.9%). This percentage increased for all pairs of stimuli in the post-training session, together with a significant increase in MI-RS values (p<0.001 for each pair of stimuli) (**Supp. Fig. 5 a-d**). In summary, MI-RS is significantly lower pre-training than in learning stage 1 (p<0.001, Mann-Whitney test), and increases across learning stages 1 to 3, before declining post-training to levels below stage 3, but above those seen pre-training (p<0.001, Mann-Whitney test, only neurons with significant MI-RS were considered; **Fig. 6d**). To obtain a direct measure of this MI-RS change for individual cells, we considered 849 neurons tracked between the pre-training and post-training imaging sessions for the three pairs of stimuli. In these neurons MI-RS increased for all pairs of object locations (KW test, p=0.987; **Fig. 6e**). This post-training effect can also be seen by a stronger contribution of non-discriminative neurons to improving decMI-RS for individual discriminative neurons (**Fig. 6f** and **Supp. Fig. 5 e-h**). These results demonstrate that learning-related changes in information generalize to stimuli not used in the task and persist even when the animal is no longer engaged in the task.

## DISCUSSION

This study provides a quantitative description of stimulus and choice information and their interplay over the course of task learning in L2/3 neurons of vS1 in mice. We find that the amount of choice information encoded by the neuronal population is more strongly tied to behavioral performance than the amount of stimulus information. Furthermore, we show that choice information is increasingly supported by a population code across learning more than stimulus information. Finally, we present data in support of our hypothesis that the emergence of choice information in vS1 reflects a more efficient use of stimulus information, resulting in changes to behavioral performance over the course of learning.

### Learning-related changes to stimulus and choice information in L2/3

The physiological manifestation of the perceptual changes observed in learning remains a focus of intense study. Previous reports have shown that neurons in the rodent vS1 and other primary areas not only carry sensory information, but can also encode multiple task variables, from navigational signals^32^ to behavioral choice^33–38^ and expectation^39^. Such representations may become stronger as animals learn behavioral tasks. We confirmed that both stimulus and choice information build up progressively during learning, with choice information being more dependent on task engagement than stimulus information. Traditionally, sensory learning has been considered as the result of an improvement in the representation of sensory inputs in primary cortex. On the other hand, recent studies have found that perceptual improvements over the course of learning may correspond to an increasingly efficient readout of sensory information in higher cortical regions while sensory representations remain stable in primary areas^27^. In our study, comparisons between the levels of mutual information and intersection information over the course of learning revealed that not only stimulus information increases but also that stimulus information is more efficiently used by neurons in primary sensory cortex during the late phase of training. In other words, stimulus and choice information do not simply increase independently of one another during task learning. Instead, the increase in readout efficacy of the stimulus information leads to the increase in choice information and, consequently, in behavioral performance. Our findings give support to both hypotheses on learning-related neuronal changes. Critically, we revealed that neurons in early sensory cortex were key in such a process, even before the involvement of higher cortical regions. Our results, therefore, are in line with a model where cortical areas are not hierarchically organized, but rather operate in parallel^38^.

In our study, we use linear decoders and information theory to directly quantify information contained in neuronal activity. Our decoder analysis shows levels of stimulus and choice decoding in vS1 that are comparable to recent reports^5,40,41^. Most studies quantified neuronal representations of information using a number of other measures, including the magnitude and frequency of neuronal activity or classification model accuracy. This difference in approach may account for some diverging observations: 1. We find that the amount of stimulus information and the number of neurons carrying it increase steadily with task training. This aligns with some previous studies^3,7,9^, but contrasts with other reports showing that stimulus-related neuronal representations remain unchanged with learning^5,11^. 2. We find that a large number of neurons carry significant levels of both stimulus and choice information. This significantly expands on previous work which segregated neurons based on stimulus and choice representation^8,34,35^. Furthermore, the level of stimulus information used to inform choice in expert mice is similar to that recently reported in A1^37^.

### Role of single neuron *versus* population codes

The relative contribution of changes in single neurons *vs*. the population to successful task learning is still unknown. Learning has been shown to change single neuron response patterns in vS1^3,9^ and elsewhere^34,42,43^, but also to influence population encoding^44^. By combining information theory with linear classifiers, we show that choice, but not stimulus, information benefits increasingly from a population code across learning. Together with the observation that stimulus and choice information emerge and are lost over different time courses in vS1, these findings indicate that different cellular and molecular mechanisms may support stimulus and choice encoding in primary sensory cortices. Furthermore, we find that when task contingencies were switched, significant changes in stimulus information are seen over the course of hours, suggesting a role for local, long-term plasticity^45,46^. In comparison, changes in choice information occur much more rapidly than would be expected if they were supported solely by local long-term plasticity, suggesting that they may be driven by an instructive top-down signal. Such signals from secondary somatosensory cortex^4^ or orbitofrontal cortex^44^ have been shown previously to be required for choice coding. In agreement with findings that these top-down signals preferentially synapse with superficial L2/3 neurons, we show that during learning choice information increases most in superficial L2. In contrast, stimulus information increases most on L3, as also seen in ^47^.

### Persistence of learning-related changes outside of task conditions

Lastly, we show that following learning, when mice are re-exposed to the same stimuli outside of the context of the task, the changes in stimulus encoding observed during learning appear persist in vS1 in a weaker, but more generalized way. Stimulus information about the task-relevant pole positions, and also nearby pole locations, increases relative to before learning began. Furthermore, it is more dependent on a population code than it was before training. This is consistent with^48^ who found that experience-induced plasticity in vS1 increased responsiveness particularly in neurons that initially showed weak stimulus responses. Together, these findings suggest that, outside of task conditions, vS1 may rely on a strengthened population code, instead of strong individual neuron responses, to continue to efficiently encode behaviorally relevant stimuli. Since this change in encoding is also context-dependent, it points to another way in which instructive top-down signals may shape how information is encoded in the vS1 population.

### Conclusion

Tools from information theory provided us with novel insights into how different types of information are encoded and integrated during learning. This approach should be of great importance in identifying promising targets for manipulation to test the causal relationship between neuronal information and behavioral performance on a related task^29^. While a growing body of work demonstrates that the manipulation of a few dozens of cortical neurons is sufficient to modulate behavior in sensory-guided tasks^8,49–51^, it remains unclear why targeting so few neurons has such an effect. Our work suggests that a common feature of such neurons could be that they carry sensory information used to inform choice, offering concrete future avenues for cracking the neural code.

## METHODS

### Subjects

All animal experimental procedures were approved and conducted in accordance with the United Kingdom Animals (Scientific Procedures) Act 1986 under project license P8E8BBDAD and personal licenses from the Home Office. Mice were housed in groups in a climate-controlled vivarium (lights on 7:00 to 19:00). The holding room temperature was 23±1 degrees Celsius and humidity was set to 40±10%. The experiments were conducted during the light portion of the photoperiod. Mice had *ad libitum* access to food, but access to water was restricted from one week before the start of behavioral training until the end of the training period. All weights were kept at 85–90% of the free-drinking weight for the duration of the behavioral experiments. All mice belonged to a GCaMP6s reporter line obtained by mating the TRE-GCaMP6s line (Jackson Laboratories strain # 024742) with the CaMKII-tTA line (Jackson Laboratories strain # 003010).

### Surgery

Eight males aged 9 to 12 weeks underwent surgery for headbar and chronic optical window implantation. Before surgery, mice received injections of meloxicam (5 mg/kg, Metacam, Boehringer Ingelheim International GmbH, Ingelheim am Rhein, Germany) and vetergesic (0.1 mg/kg, Ceva Animal Health Ltd, Amersham, UK). They also received a marcaine (AstraZeneca, Cambridge, UK) injection under the scalp. Eye cream was applied to the eyes (lacri-Lube, Allergan, UK). Anesthesia was induced via inhalation of 4% isoflurane (Zoetis, Leatherhead, UK) at 1 L/min. When mice were fully anesthetized, they were placed in a stereotaxic frame (Kopf instruments, Tujunga, CA). Depth of anesthesia was monitored by checking pedal withdrawal reflex and respiration rate. Body temperature was kept at 37±1°C. Isoflurane rate was kept at 0.8–1.2% at 0.7 L/min during surgery. A circular incision was made into the scalp, the skull was cleaned, and the periosteum removed. A 3 mm diameter craniotomy was centered over the right vS1 following stereotaxic coordinates (3.1 mm lateral from the midline and 1.3 mm posterior from the Bregma suture). The dura mater was left intact. The craniotomy was then sealed with two glass coverslips (3 mm and 4 mm diameter, Thermo Fisher Scientific, UK) glued to one another using optical adhesive (Norland, New Jersey, USA). A stainless steel headbar was cemented onto the skull using dental cement (Super-Bond C&B, Sund Medical, Japan). After surgery, mice were allowed to recover for one to two weeks before starting handling and water regulation. Handling and gentle restraint by the experimenter were performed over three days. Mice were then habituated to be headfixed under the imaging setup, and to receive water from a spout placed in front of them. This habituation phase lasted three further days, after which behavioral training started.

### Sensory stimulation

Tactile stimuli consisted of a small metallic pole (a blunt 18G needle, diameter 1.27 mm), held vertically and contacting the majority of the left whiskers of the mouse for 1 to 1.5 seconds at approximately 0.3 cm from the whisker pad. Mice were free to whisk against the pole. The pole was connected to a perpendicular plastic arm mounted onto the shaft of a stepper motor (RS PRO Hybrid 535-0467; RS Components, UK). The stepper motor was mounted onto a motorized linear stage (DDSM100/M; Thorlabs. controlled by a K-Cube Brushless DC Servo Driver [KBD101; Thorlabs]), which moved the metallic pole close to the whiskers or away from them. The length traveled by the linear stage was identical during Go and No-go trials. During the pre-training and post-training sessions, the pole contacted the whiskers in six positions along the antero-posterior axis of the animal, separated 2.4 mm from one another. The most anterior position is denoted as position 1 throughout the text, while the most posterior is position 6. During the behavioral training phase, positions 3 and 6 were the only two used as tactile stimuli. The stepper motor only rotated between the positions once it had traveled away from the whisker pad via the linear stage. Rotating from position 1 into position 6 took approximately 90 ms longer than rotating into position 3. The sound frequency emitted consisted primarily of energy below 1 kHz, which is outside the mouse frequency hearing range^*52*^. Sound intensity of the stepper motor and linear stage was <30 dB SPL. Ambient noise inside the microscope box was below 40 dB SPL. Intensity thresholds for primary auditory cortical neurons in the mouse range between 4 and 39 db SPL^*53*^. Our measurements allowed us to exclude the presence of potential auditory cues during the task.

### Behavioral training

Hardware and software for behavioral experiments were controlled through the open-source toolbox pyControl (OEPS Electrónica e Produção, Alges, Portugal)^54^. We trained mice on a whisker-based object localization Go/No-go task^55^. As described in the previous section, the metallic pole came into contact with the whiskers for 1 to 1.5 s in one of two possible positions along the anterior-posterior axis of the mouse. The first lick latency was calculated from stimulus offset. During Go trials, mice were rewarded with an 8 μl drop of sweetened water (10% sucrose solution) when they licked from a spout during a response / licking window starting 100 ms after the retraction of the pole and lasting four seconds (Hit trials). Water was not delivered if mice licked while the pole was still in contact with the whiskers (“Too soon” trials, not included in the analysis). Licks during No-go trials were considered as False Alarms and were punished with an extended inter-trial interval (time-out). No punishment nor time-out were presented when mice did not lick during Go trials (Misses). Daily training took place in three consecutive blocks of about 16 minutes duration each. Across the three blocks, mice performed on average 187±48 trials per training day. Learning was classified into three stages, based on the percentage of correct responses in each block: 0-55% (stage 1), 55-75% (stage 2), 75-100% (stage 3). Training ended when a mouse’s performance averaged higher than 70% across the three blocks, for three consecutive days. Only mice that performed above 70% for three consecutive days were retained for analysis (n=8). During pre-training and post-training sessions, mice (n=3) were not water-regulated but had *ad libitum* access to water in their home cage. This was to avoid potential confounds related to the animals being thirsty during these sessions. Two of the mice imaged pre- and post-training were also used in “switch” session. At the end of the switch session, each of them was tested on the task based on the original contingencies. They both reached 70% within one session, at the end of which training was ended.

### Two-photon imaging

Quasi-simultaneous double-plane two photon calcium imaging was performed using the set up described in detail in^17^. Two photon excitation light was emitted by a femtosecond Ti:Sapphire laser (MaiTai BB, Spectra Physics, USA) tuned to 900 nm. Double-plane imaging was achieved using a system including a DTSX-400-980 Acousto-Optic Deflector (AOD; Photon Lines Ltd., UK), a SF11 equilateral prism (Thorlabs, UK), and two aspheric lenses (C330TMD-B, Thorlabs, UK). The laser beam was directed to the AOD, whose acoustic frequency switched between 83.5 MHz and 91.5 MHz, generating two optical paths: one leading to the nominal focal plane, and the second encompassing the aspheric lenses for refocusing onto a second focal plane, placed 130 μm below the first one. This permitted efficient acquisition of multiple planes while keeping the behavioral experiment short. During each of the three behavioral blocks (ca. 16 min, see above), we re-focused the imaging path once to acquire imaging data from fields of view (FOVs) at four depths below the cortical surface (−130, -190, -260 and -320 μm), equivalent to approximately cortical layers 2 and 3. The same FOVs were imaged in each mouse across training sessions. The two beams were then recombined through a polarizing beamsplitter. Calcium transients were acquired using a Sutter Moveable Objective microscope (MOM, Sutter, USA) controlled by ScanImage 5.2.1 software (http://scanimage.org) with minor modifications for the AOD beam steering control. The beam was scanned through an 8 kHz resonant scanner in the x-plane and a galvanometric scanning mirror in the y-plane. The resonant scanner was used in bidirectional mode, at a resolution of 512 × 512 pixels, allowing us to acquire frames at a rate of ∼15 Hz per imaging plane. A 16X/0.80W LWD immersion objective (Nikon, UK) was used. Laser power, as measured under the microscope objective, was between 80 mW and 95 mW. Emitted photons were guided through a 525/50 filter onto GaAsP photomultipliers (Hamamatsu Photonics, Japan). Neuronal fields were 400 × 400 μm in size.

### Intrinsic Optical Signal Imaging

Intrinsic Optical Signal Imaging (IOSI) was carried out at the end of the experimental procedure to confirm that 2p imaging was performed in vS1. General anesthesia was induced with 4% isoflurane (Zoetis, Leatherhead, UK) at 1 L/min, and was then kept at 0.6–0.8% at 0.7 L/min during imaging. An *intra muscular* injection of chlorprothixene hydrochloride (1 mg/kg) was administered to inhibit whisker movements. Mice were head-fixed and placed on a heated mat. Temperature was kept at 37±1°C. One whisker from the left row B, C or D was identified and threaded through a glass capillary, which was attached to a ceramic piezoelectric stimulator (*e*.*g*., PB4NB2W Piezoelectric Bimorph Bending Actuator with Wires, Thorlabs). If the surrounding whiskers touched the external side of the capillary, they were carefully trimmed using a pair of iris scissors under a dissecting microscope. A Retiga R1 camera with a 50 mm and a 135 mm lens (Nikon) attached in tandem configuration was used for imaging^56^. Imaging was performed through the chronic cranial window previously implanted over the right parietal lobe. The whisker stimulation protocol consisted of 1 s stimulation at 10 Hz with 20 s ITI, repeated 40 times, for a total of 400 deflections. This protocol was repeated 3-4 times per mouse on different whiskers in order to map the barrel fields in vS1. For post-hoc confirmation of the imaging location in vS1, a map of the vS1 barrels obtained through IOSI was overlaid upon images of the areas investigated with 2p imaging (see, **Fig. 1**).

### Two photon imaging analysis

Raw 2p images were imported into the Suite2p software (https://github.com/MouseLand/suite2p^57^), which performed correction for mechanical drift along the x and y axes, image segmentation, and neuronal and neuropil trace extraction. We manually inspected all regions of interest detected by the Suite2p built-in classifier, to confirm that they corresponded to neurons rather than structures such as fragments of neuronal projections or perpendicular blood vessels. For each confirmed neuron, the signal at each time frame (F***(t)***) was calculated as the average fluorescence of all pixels inside the ROI. The time series of the neuronal calcium trace and the neuropil calcium trace were exported to Matlab for further analysis. Baseline Fluorescence (F_*0*_) was considered to be the median of the 10th to the 70th percentile of the fluorescence distribution across all frames acquired. Each neuron’s fluorescent trace was then corrected for the baseline using the formula: (F(t) – F_0_)/F_0_, commonly denoted as ΔF/F_0_. Subtraction of the neuropil signal was applied to each neuron’s trace as described previously^58^, using a contamination ratio of r = 0.7. Semi-automatic ROI registration across the pre-training and post-training imaging sessions was performed using the “registers2p” package (https://github.com/cortex-lab/Suite2P/tree/master/registers2p). Although the majority of our analysis did not require to systematically track neurons over the course of daily imaging sessions, examples of tracked neurons are given, *e*.*g*., in Fig. 1g or Fig. 6e. The signal-to-noise ratio (SNR) was calculated during pre-training and post-training imaging sessions on the raw fluorescent trace for each ROI. The signal was the maximum fluorescence value of the whole trace. The noise was the standard deviation of the distribution of fluorescence values recorded during the first 3 seconds of acquisition, when sensory stimulation was not yet present^17^.

### Mutual information analyses

The information content of the neuronal response about stimulus (MI-RS) and choice (MI-RC) has been quantified using Shannon’s Mutual Information. Mutual information between two discrete random variables is defined as:

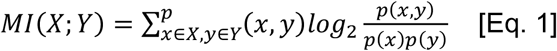

We used a plug-in direct method in which probabilities are computed directly from discretized data and then plugged into the information equation, as follows. Continuous values of the ΔF/F_0_ traces were discretized using two equally populated bins. Calculation of the marginal and joint probabilities in Eq. 1 was performed by building a histogram of the discretized values of the neuronal response and stimulus/choice across trials. Values of MI obtained were corrected for limited sampling bias using the Panzeri-Treves method^59^. For each time frame during the trial (3 seconds after stimulus onset), a single calculation of MI was performed by considering the neural response as the set of values of ΔF/F_0_ for that time point across trials. Binning of the ΔF/F_0_ values has been performed independently at each time frame. Assessment of significance for mutual information values was performed through a permutation test limited to the first second of neural activity after stimulus onset (i.e., before the opening of the time window for licking). A null distribution of MI values was built through randomly permuting the neuronal response across trials^60^. Permutation of trials was performed consistently across all time points. This allowed us to abolish the association between stimulus (or choice) and neuronal response while preserving the autocorrelation in the fluorescence signal. Evaluation of the significance level of the MI carried by each neuron was performed as follows. For each time point, 50 bootstrapped values of MI were calculated. Bootstrapping was performed without correcting for limited sampling bias, owing to the larger statistical power of non bias-corrected null distribution^61^. Significance of MI for a single neuron was assessed by comparing (using the Kruskal-Wallis test) the distribution of non bias-corrected MI across frames, with the distribution of the mean values of bootstrapped MI for the same frames. Values of MI reported for significant neurons are corrected for limited sampling bias. Within a trial, MI-RC peaked about 1 second after the pole moved away from the whiskers (**Fig. 2g**). This peak comes later than the mean first lick latency in Hit trials observed during stage 2 and 3 of learning but is in line with the first lick latency during FA trials (**Fig. 1d**). However, since mice do not lick during CR and Miss trials, a behavioral paradigm different from the Go/No-go used here would be required to properly evaluate any temporal coincidence between choice and peak MI-RC.

### Intersection information analyses

Intersection Information^28,29^ is an information-theoretic measure quantifying the amount of stimulus information, present in the neural activity, that is used to inform choice. The quantity is based on the Partial Information Decomposition formalism^62^, and is bound by both MI-RS and MI-RC. This property allows one to conveniently express II/MI-RS as the fraction of stimulus information that is used to inform choice. We computed II following the procedure in ^28^ and using the same plug-in direct method used for MI-RS and MI-RC and described above. Correction of limited sampling bias has been performed using quadratic extrapolation^63^. To reduce the computational cost of the analysis, the dimensionality of the ΔF/F_0_ traces in the post-stimulus epoch was halved by considering the average ΔF/F_0_ across two consecutive time frames. Discretization of the neuronal response was performed, independently at each time frame, using two equally populated bins. Assessment of the significance of II for a single neuron was performed using a permutation test, similarly to what we did for mutual information. The null distribution for II, at each time frame, was calculated by permuting neuronal responses across all trials.

### Decoder analyses

Linear logistic regressions were trained using 5-fold cross-validation with an L_2_ penalty. Decoders were trained separately on each session FOV, *i*.*e*., each FOV recorded during each imaging session. All decoders received, as input, neuronal activity from the first second following stimulus onset, and all trial types except ones where mice licked too soon were included (Hit, FA, CR, Miss). This ensured that the inputs to the decoders did not coincide with any licking, creating confounds. Decoders trained on individual neurons received as input the ΔF/F_0_ response of a single neuron (**Fig. 3a-b, Supp. Fig. 2** and **Supp. Fig. 4e**), whereas decoders trained on multiple neurons received as input the concatenated ΔF/F_0_ responses of all included neurons (**Fig. 3c-j, Fig. 4a-b, Fig. 6f**, and **Supp. Fig. 4f-h**).

Decoders were trained to predict for each trial either stimulus identity (Go/No-go) (**Fig. 3a & c & e-g, Fig. 6f**, and **Supp. Fig. 4e-h**) or choice (Lick/No-lick) (**Fig. 3b & d & h-j**), or both (**Fig. 4a-b**). To evaluate decoder performance, we computed a mutual information score on the confusion matrices obtained from the validation fold predictions (decMI)^22^. This approach allows us to estimate information content not only for individual neurons, but also for groups of neurons. We confirmed the validity of our decMI measure by comparing decMI values obtained for individual neurons to the previously computed MI measures (**Supp. Fig. 2**). Decoder performance was reported as decMI-RS for stimulus identity (MI when predicting stimulus from neural response) and decMI-RC for choice (MI when predicting choice from neural response). We report results grouped across depths, at each stage of learning.

When decoder performance for individual neurons was plotted against behavioral performance, jitter was added to the behavioral performance for visualization purposes to enable values for individual neurons to be distinguished. This was done by resampling the behavioral performance value for each neuron using a normal distribution centered on the true value with a standard deviation of 1.2% (**Fig. 4a-b**). For all decoders, to ensure sufficient numbers of each trial type were available for training the decoders, for choice decoding, only sessions with at least six Lick and six No-lick trials were included (number of session FOVs removed: eight for stage 1, on for stage 2, zero for stage 3) (**Fig. 3a & c & e-g, Fig. 4a-b** and **Supp Fig. 2b**). Furthermore, to ensure that choice decoding and stimulus decoding results could be appropriately distinguished, only sessions that included at least six correct and six wrong trials were included for all decoders trained on task training sessions (number of session FOVs removed: zero for stage 1, four for stage 2, 23 for stage 3) (**Fig. 3, Fig. 4a-b** and **Supp Fig. 2**).

Null distributions were estimated by running, for each neuron, ten 5-fold cross-validations, each computed on data where the trial types had been randomly shuffled. By aggregating decMI values resulting from shuffled data across neurons from the same session FOV, a null distribution over decoder performance was constructed for each session FOV. Neurons with decMIs above the 95^th^ percentile of the null distribution were identified as carrying significant information and labeled as “discriminative” neurons for the session FOV. Neurons that did not meet the threshold for their session FOV were labeled as “non-discriminative” (**Fig. 3a & c** and **Supp Fig. 4e**).

To compare decMI and MI values for each neuron, we recomputed both maximum MI and MI significance over the same trial length used for the decoders, i.e., the first second following stimulus onset. The MI-RS values thus obtained were then compared to decMI-RS for each neuron (**Supp. Fig. 2a**), whereas MI-RC values were compared to decMI-RC values (**Supp. Fig. 2b**). For each stage of learning, a linear regression model was fit to the data, and the goodness-of-fit was measured using the R^2^ coefficient of determination (**Supp. Fig. 2a**). The same process was repeated to compare MI-RC and decMI-RC values for neurons significant for both (**Supp. Fig. 2b**).

To evaluate how decoder performance changed as neurons were added to the pool of data provided to the decoders, neurons were first ordered from the lowest to highest individual decMI(R;S) (**Fig. 3c** and **Supp. Fig. 4f**) or decMI(R;C) (**Fig. 3d**). Decoders were trained to classify stimulus or choice, respectively, as neurons were added in that order, for each FOV from either the full pool of neurons (**Fig. 3c & d**, left and **Supp. Fig. 4f**, left) or the pool of non-discriminative neurons (**Fig. 3c & d**, right and **Supp. Fig. 4f**, right). Since different FOVs comprise different total numbers of neurons, to enable the pooling, the data from each FOVs was downsampled down to the lowest number of neurons present in at least 5 FOVs (18 neurons for the full population, and 11 neurons for the non-discriminative pool). Neuron numbers were then converted to percentages and mean ± sem was calculated across FOVs. Statistical comparisons between stage 1 and 2, as well as stage 2 and 3 were computed for each datapoint using t-tests. As elsewhere, p-values were Bonferroni corrected for multiple comparisons, calculated here as the total number of comparisons for each data panel.

To determine the gain contributed by non-discriminative neurons to classification performance for discriminative neurons, in addition to training decoders on data from individual discriminative neurons, additional decoders were trained on data from each individual discriminative neuron paired with all the non-discriminative ones from the same session FOV. These decoders received as input the concatenated responses of all included neurons for each trial. The gain contributed by the non-discriminative population was then measured by, for each discriminative neuron, subtracting the decMI of the decoder trained only on the individual discriminative neuron data from the decMI of the decoder trained on data from the same discriminative neuron and the pool of non-discriminative neurons (**Fig. 3e & h** and **Supp Fig. 4g**). Positive gain reflected an improvement when including non-discriminative data, whereas negative gain reflected a drop in performance. Histograms over these gains were computed by binning the data into 40 bins (**Fig. 3f & i** and **Supp Fig. 4h**). Cumulative distributions over the same data, binned into 80 bins (**Fig. 3g & j** and **Fig. 6f**).

Lastly, to determine whether decoders trained on stimulus classification performance carried choice information, decoders were first trained on the entire pool of neurons for each session FOV to classify stimulus identity (Go vs NoGo). We then sorted trials according to whether the decoder classified stimulus identity correctly (“S+”, *e*.*g*., for a trial where the stimulus was classified as Go and it was Go) or incorrectly (“S-”, *e*.*g*., for a trial where the stimulus was classified as Go but it was NoGo).

Finally, we calculated the % correct choice across session FOVs for each stage of learning enabling corresponding animal behavior (% correct choice in S+ trials - % correct choice in S-trials). Paired t-tests were then computed between both behavioral performances in order to determine whether the performance levels observed differed significantly at any learning stage for S+ *vs*. S-trials.

Although decoder mutual information is primarily reported in this paper, we also computed balanced accuracies for the decoders (data not shown) and found that the decoding accuracies computed for discriminative neurons at all stages were comparable to those reported in previous work on decoding tactile stimuli and choice from neural activity^3,11,41^.

### Statistical analysis

Differences between the distributions over population data for pairs of learning stages were evaluated using two-sided Kolmogorov-Smirnov tests with Bonferroni corrections for multiple comparisons. Corrected p-values <0.05 are considered significant. Mean ± standard error of the mean (sem) is reported unless otherwise indicated. One, two and three significance stars indicate p <0.05, p <0.01, and p <0.001 respectively.

### Analysis software

Mutual Information calculations were carried out using the Information Breakdown Toolbox^64^. Intersection information calculations were performed using custom MATLAB routines, coupled with Python package BROJA2-PID^65^. Decoder analyses were performed in Python 3.9 with custom scripts developed using the following packages: NumPy^66^, SciPy 1.6.2^67^, Pandas^68^, Matplotlib^69^, and Scikit-learn 0.24.1^70^. The remaining analysis was using custom scripts written in MATLAB2021b.

## ACKNOWLEDGEMENTS

We thank Dr Ana Bottura de Barros and Dr Severin Limal for help with the behavioral set up, Dr James Rowland for help on the preprocessing of 2-photon imaging data, Dr Liad Baruchin and Dr Severin Limal for help with Intrinsic Optical Signal Imaging.

This work was enabled by the resources provided by Compute Ontario and the Digital Research Alliance of Canada (www.computeontario.ca and www.alliancecan.ca).

## FUNDING

This work was supported by the Wellcome Trust (109908/Z/15/Z, to M.M.K.) and the Human Frontiers Science Programme (RGY0073/2015, to B.A.R. and M.M.K.). M.P. and R.M. are Marie Sklodowska-Curie Fellows (EnlightenedLoom - 101024523, and MoWS - 894032). C.J.G. was supported by an NSERC Canada Graduate Scholarship - Doctoral Program, and an Ontario Graduate Scholarship. S.P. was supported by EU H2020 under Grant Agreement No. 945539 (Human Brain Project SGA3). B.A.R. was supported by a CIFAR Catalyst grant, a CIFAR AI Chair grant, an NSERC Discovery grant (RGPIN-2014-04947), and an Ontario Early Researcher Award (ER17-13-242).

## SUPPLEMENTARY INFORMATION

### Expanded data and statistics

#### Population decoding

The correlation between stimulus MI measures (decMI(R;S) *vs*. max MI(R;S)) was strong for all stages, with a slope of nearly 1, pointing to a near-identity relationship. In addition, most neurons had consistent significance statuses, being significant either for both measures (MIRS+decMIRS+) or for neither (MIRS-decMIRS-) (77.5±0.8% in stage 1, 71.6±1.4% in stage 2, 73.2±1.4% in stage 3). For choice MI measures, the correlation between decMI(R;S) and max MI(R;S)) was initially weak in stage 1, increasing substantially and reaching a near-identity relationship by stage 3. Nonetheless, most neurons achieved consistent significance statuses with both measures (80.6±1.2% in stage 1, 70.1±1.1% in stage 2, 73.2±1.7% in stage 3). Together, these findings support the use of decMI as an estimate of information content in neural activity (**Supp. Fig. 2**).

We found that as mice improved in their task performance, median decMI-RS remained within the null distribution range (stage 1: 0.09, stage 2: 0.10, stage 3: 0.09). However, the percentage of discriminative neurons, *i*.*e*., neurons performing above the null distribution range (p<0.05), increased 1.5-fold from stage 1 to stage 3 (stage 1: 16.0±1.2%, stage 2: 19.5±1.1%, stage 3: 24.4±2.2%, mean±sem across FOVs; **Fig. 3a**). Although many neurons still showed chance-level decoding decMI-RS in stage 3 of learning, the overall distribution across neurons changed from stage 1 compared to stages 2 and 3 (ks-test p=0.013 for stage 1 *vs*. stage 2, p=0.005 for stage 1 *vs*. stage 3, but p>0.05 for stage 2 *vs*. stage 3), with the 95^th^ percentile of the decMI-RS distribution across neurons increasing with each stage (0.31 in stage 1, 0.39 in stage 2, 0.47 in stage 3). Similarly, the decMI-RC distribution changed across learning, with the 95^th^ percentile going from 0.21 in stage 1 to 0.25 in stage 2, and 0.32 by stage 3 (ks-test p=0.005 for stage 1 *vs*. stage 2, p<0.001 for stage 1 *vs*. stage 3, p=0.019 for stage 2 *vs*. stage 3; **Fig. 3b**). Again, although most neurons still showed chance-level decMI-RC by stage 3 (median decMI-RC: 0.09 for all stages), the percentage of discriminative neurons also rose consistently, more than doubling from stage 1 to 3 (7.4±1.0% in stage 1, 11.9±0.8% in stage 2, 19.6±2.0% in stage 3). When considering the information gain via non-discriminative neurons, for decMI-RS the 5^th^ percentile of the gain stayed around -0.21, and the 95^th^ percentile rose only slightly (stage 1: 0.32, stage 2: 0.34, stage 3: 0.37), and only stages 2 and 3 differed significantly from one another (ks-test p=0.006 for stage 2 *vs*. 3; **Fig 3f**). For decMI-RC the 5th percentile increased only slightly (stage 1: -0.26 in stage 1, stage 2: -0.21, stage 3: -0.19; **Fig 3i**).

Decoder analysis was also used to test the hypothesis that, as mice learn to perform the object localization task, sensory information should increasingly inform the animal’s choice. We trained linear decoders to classify Go *vs*. No-go stimuli from the responses of the full neural population for each FOV. We then calculated whether the decoder classified the stimuli correctly or incorrectly. Next, we quantified the behavioral impact of the stimulus encoded in neural populations by computing the difference in behavioral performance between trials where the stimulus was classified correctly or incorrectly by the decoder. If the sensory information encoded in neural activity is not used to inform behavioral choices, then there would be no difference in behavioral performance between correctly decoded trials (trials in which neural activity provides faithful stimulus information) and incorrectly decoded trails (where neural activity gives misleading stimulus information). In stage 1, despite the presence of stimulus information, there was no difference in performance between correctly and incorrectly decoded trials, meaning that stimulus information is present but not effectively used. In contrast, in stage 3, mice performed significantly better in correctly classified trials, compared to incorrectly classified ones (paired t-test p>0.05 for stages 1 and 2, p=0.029 with Cohen’s D of 0.69 for stage 3; **Supplementary Fig. 3a & b**). These results suggest that, as mice improve in their task performance, the sensory information encoded in vS1 neural activity is more efficiently read out by downstream structures to inform behavior.

#### Intersection information

Intersection Information can be positively skewed, from stage 1 to stage 3, by the increased correlation between stimulus and choice. We finally sought to control for this effect by subsampling trials to equalize the behavioral performance at 75% across all learning stages. For each ROI, a single, independent, random subsampling of trials was performed. II and MI-RS were calculated on the subsampled trials in the same way as done for the full data. Matching the behavioral performance causes a substantial reduction of the available trials (**Supp. Fig. 4c**), especially in stage 1 and stage 3. This increases the limited sampling bias for II and MI-RS and makes it impossible to directly compare both the values of II/MI-RS as well as the number of significant neurons resulting from the full data and the subsampled data analyses. When fixing the behavioral performance, we still observed an increase in the median II/MI-RS between stage 1 and stage 3, across all depths (**Supp. Fig. 4a**). The results of this analysis also confirm an increase in the number of neurons significantly encoding II from stage 1 (−130 μm: 20.6%, -190 μm: 19.2%, -260 μm: 21.5%, -320 μm: 13.4%) to stage 3 (−130 μm: 31.0%, -190 μm: 33.1%, -260 μm: 33.9%, -320 μm: 29.3%) (**Supp. Fig. 4b**). Overall, these results confirm how the observed increase in readout efficacy of the neural code in vS1 from stage 1 to stage 2 goes beyond the trivial increase in correlation between stimulus and choice with learning.

#### Switch session

For the three mice used, individual behavioral performances on the switch day were 41.9%, 35.6% and 36.1% over 146, 157 and 163 trials, respectively. The mean MI-RS across all neurons during the switch session (n=154 neurons at -130 μm, 217 at -190 μm, 149 at -260 μm and 158 at -320 μm) was between 1.2 and 2.9-fold lower than the MI-RS in the same FOVs during stage 3 of learning (1.18 at -130 μm, 2.93 at -190 μm, 1.63 at -260 μm, 1.81 at -320 μm) (**Supp. Fig. 5a**). At the same time, MI-RS was higher at all depths than the values observed during stage 1, when the association between stimulus position and water reward had not yet been established. This effect was stronger in superficial layer 2 neurons compared to neurons at lower depths within the cortical column (stage 1/switch MI-RS = 0.217 at -130 μm, 0.451 at -190 μm, 0.743 at -260 μm, 0.528 at -320 μm). These results suggest that consistency in the pairing between a sensory stimulus and a water reward is necessary to maintain higher levels of sensory information in vS1 neurons. A similar trend was observed for choice information (MI-RC), which dropped compared to stage 3 of learning, and had values similar or even lower than those observed at stage 2 (1.8-folds lower than stage 3 at 130 μm, 2.9-folds at 190 μm, 1.9-folds at 260 μm, 2-folds at 320 μm; **Supp. Fig. 5b**).

#### Pre- and post-training

Stimulus information increased in the neurons from 2 to 3-fold after training (1.94-folds at -130 μm, 2.78 at -320 μm), as did the percentage of neurons carrying significant MI-RS (between 11.7% at 130 μm and 28.2% at 260 μm). We sought to assess what these changes from pre-to post-training reflected in terms of discriminative and non-discriminative contributions to population encoding of stimulus information. As before, decoders were trained on the binary task of differentiating trials in which the stimulus pole was in the Go position from those where it was in the No-go position, and MI was calculated on the resulting confusion matrices. We found a much more modest proportion of discriminative neurons, going from 7.2±1.1% pre-training to 10.4±1.6% post-training, with a slight shift upward in the decMI-RS distribution for the population (ks-test p=0.002; median: 0.07 pre to 0.08 post; 95th percentile: 0.21 pre to 0.23 post; **Supp. Fig. 6e**). When it came to population encoding of stimulus information, no difference between pre- and post-training decMI-RS levels were observed as neurons were added to the decoder pool, in contrast to the increase observed during training, when all neurons were included (**Supp. Fig. 6f**). However, the contribution of non-discriminative neurons to improving decMI-RS for individual discriminative neurons was stronger than observed during learning for stimulus information (**Fig. 3e-g**) with the gain distribution shifting positively (ks-test p<0.001; median: -0.09 pre to 0.01 post; 5^th^ percentile: -0.28 pre to -0.23 post; 95^th^ percentile: 0.27 pre to 0.41 post) (**Fig. 6f** and **Supp. Fig. 6g & h**). These results suggest that although the strength of stimulus information carried by individual neurons decreases when the association of the stimulus to a reward is withdrawn, the more robust encoding of stimulus information developed by the population as a whole is maintained.

## Supplementary Table

**Supplementary Table 1:**
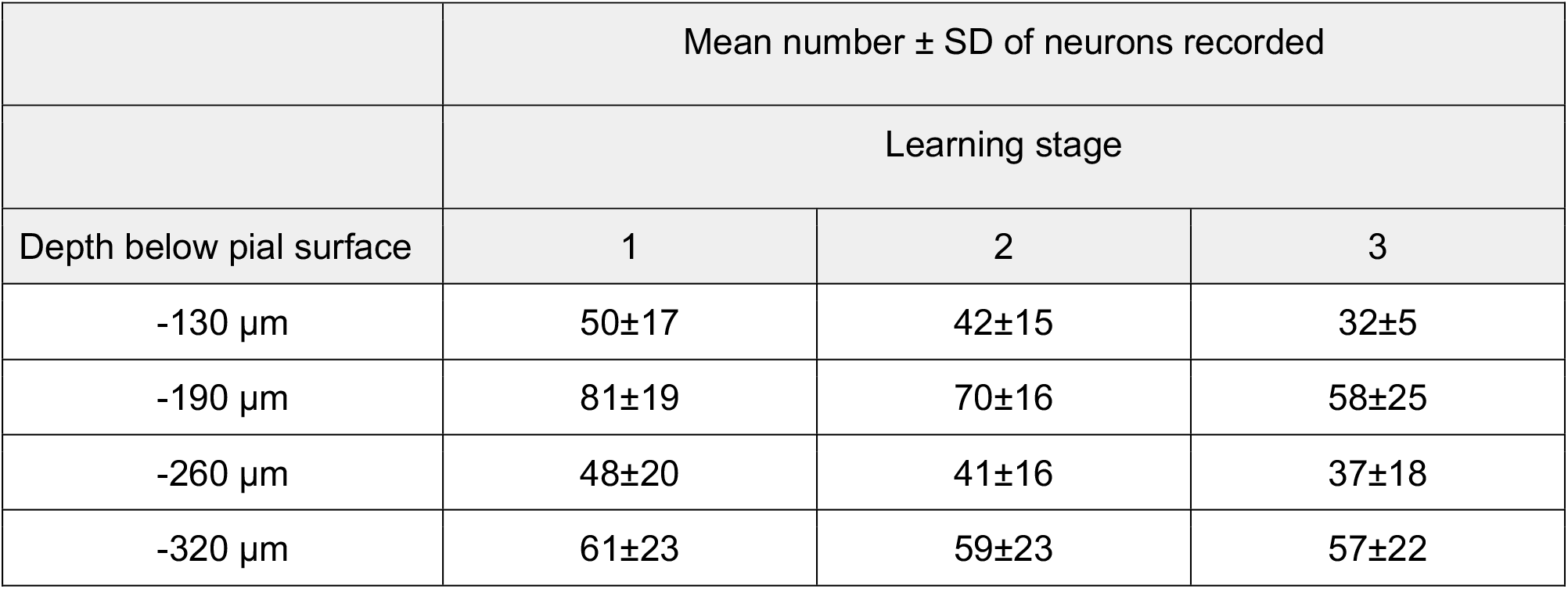
Summary of recording sessions. The number of imaged neurons decreased slightly between stage 1 and stage 3 at -130 μm and -190 μm: p<0.01 and p<0.05, respectively, Mann-Whitney test. It remained stable at -260 μm and -320 μm: p>0.05 for both depths, Mann-Whitney test; **Supp. Fig. 1**). The signal-to-noise ratio (SNR, see Methods) remained stable over time at each of the depths tested (p>0.05), indicating that imaging quality did not deteriorate over weeks. When tested across depths, SNR was comparable, except between -130 μm and -320 μm (p = 0.012, Kruskal-Wallis (KW) test corrected for multiple comparisons).

| | Mean number ± SD of neurons recorded | | |
| --- | --- | --- | --- |
|  | Learning stage |  |  |
| Depth below pial surface | 1 | 2 | 3 |
| -130 μm | 50±17 | 42±15 | 32±5 |
| -190 μm | 81±19 | 70±16 | 58±25 |
| -260 μm | 48±20 | 41±16 | 37±18 |
| -320 μm | 61±23 | 59±23 | 57±22 |

## Supplementary Figures

**Supplementary Figure 1:**
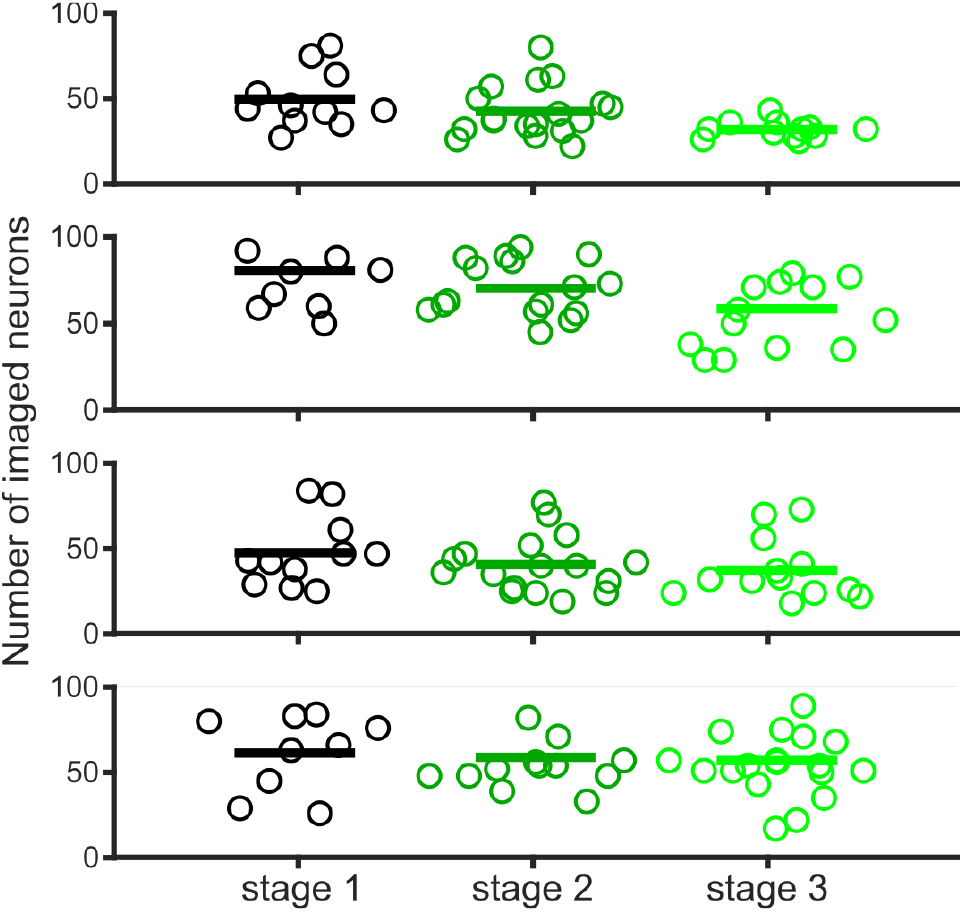
The number of imaged neurons remains stable across imaging sessions. Number of neurons imaged in FOVs at each cortical depth and each learning stage (stage 1: black, stage 2: dark green, stage 3: light green). Each circle represents a FOV. Horizontal lines indicate the mean across FOVs. See Supp. Table 1 for data.

**Supplementary Figure 2:**
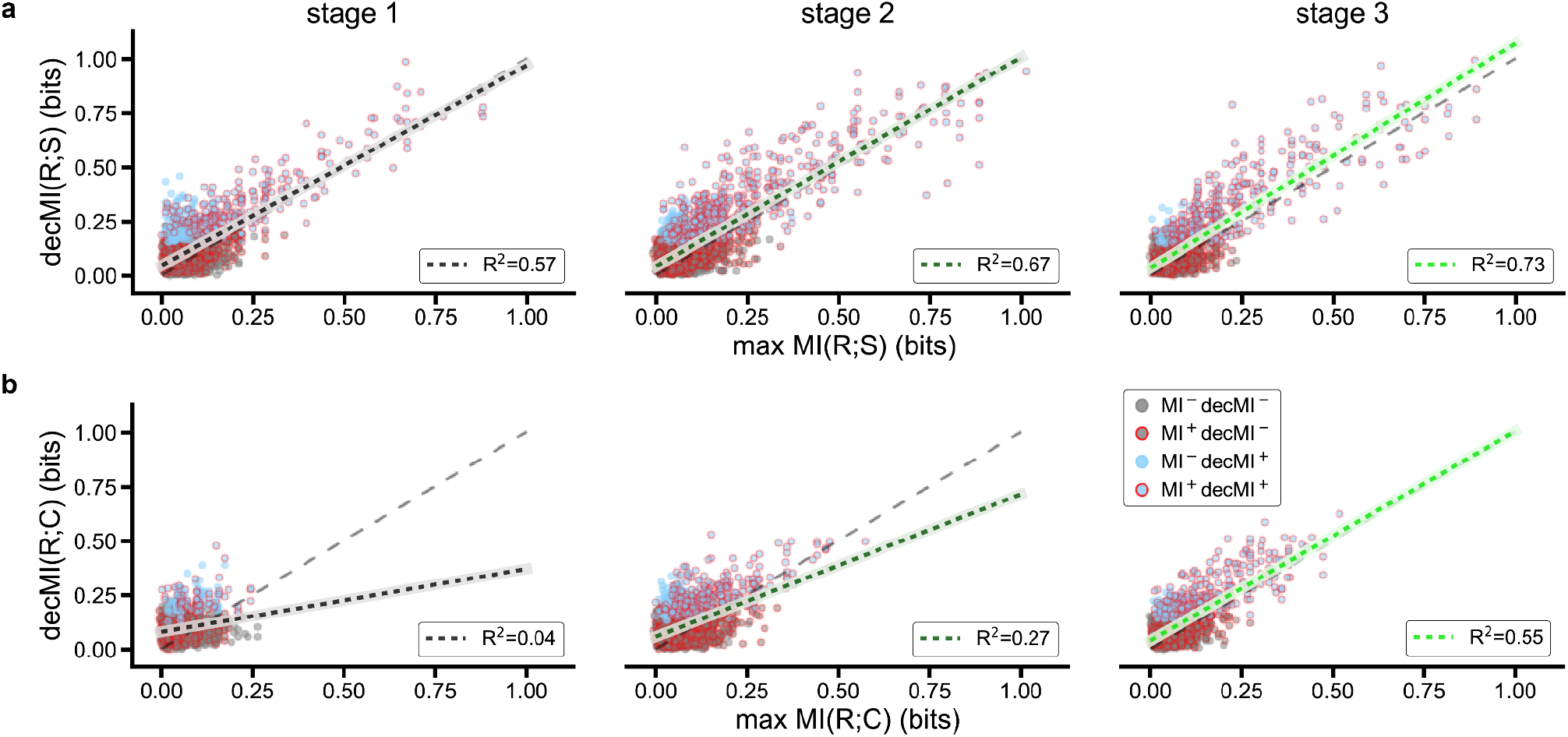
Mutual information calculated directly on neuronal responses correlates strongly with mutual information calculated from decoder performances for individual neurons. **a**. Maximum MI-RS is plotted against the MI-RS calculated from the decoder performances (decMI-RS) (both calculated over the first second following stimulus onset) for stages 1, 2 and 3 (left to right). As shown in the shared legend in b, dots are plotted in blue for discriminative neurons (decMIRS^+^), and gray otherwise (decMIRS^-^). Dots are also plotted with a red edge if MI is significant (MIRS^+^). The light dashed line is the identity line, and the correlation fit line is plotted in color for each stage (R^2^: 0.57 in stage 1, 0.67 in stage 2, 0.73 in stage 3). **d**. Same as for c, but for MI-RC *vs*. decMI-RC (choice). The correlation fit lines are plotted in the stage’s color (R^2^: 0.04 in stage 1, 0.27 in stage 2, 0.55 in stage 3).

**Supplementary Figure 3:**
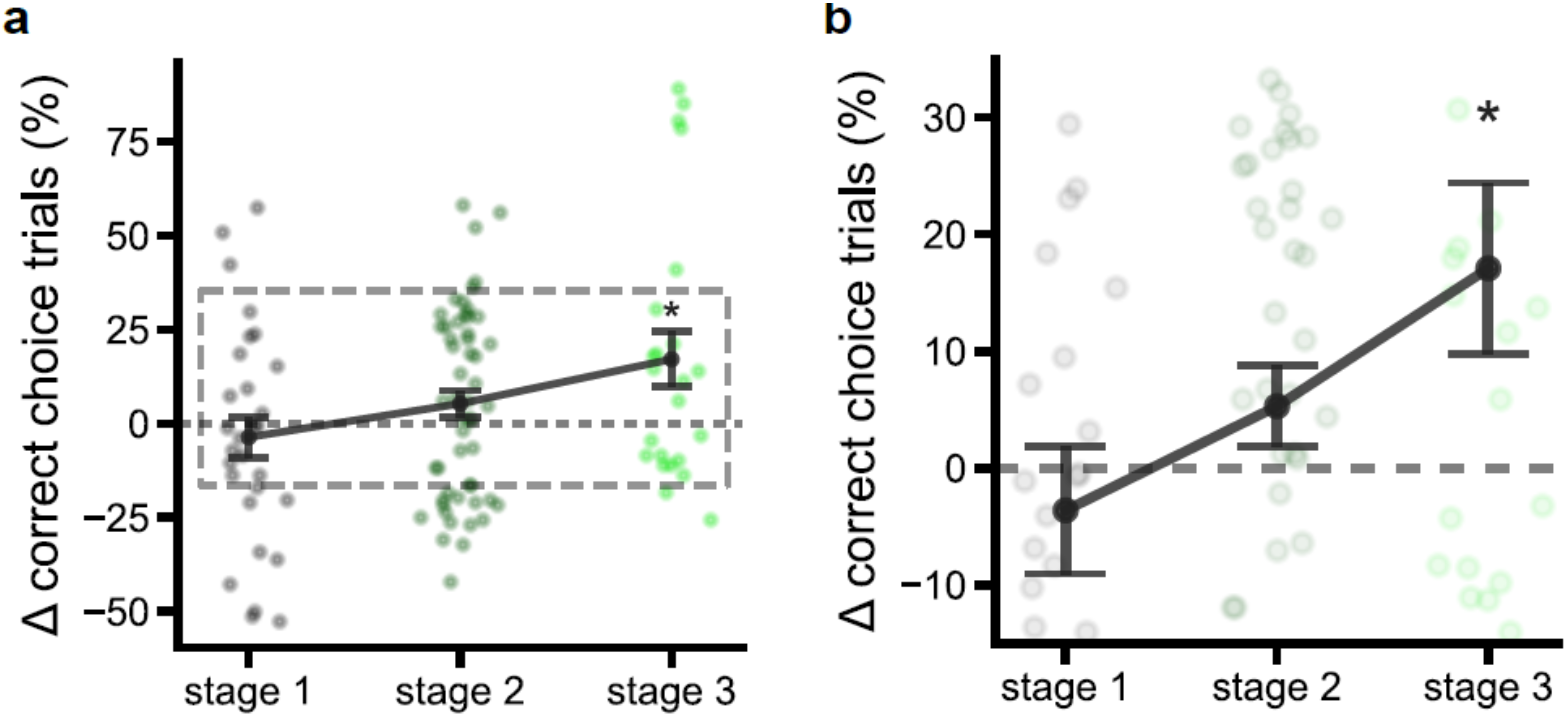
Decoder analysis confirms that stimulus information increasingly instructs behavioral choice as learning progresses. **a**. Differences in behavioral performance (% trials in which the correct choice was made) for all learning stages across trials for which the stimulus was correctly versus incorrectly classified by a decoder trained on the full population. Mean across FOVs is plotted, and dots show data for individual sessions, pooled across depths. Asterisk indicates values significantly different from 0 (paired t-test: p<0.05). **b**. Zoomed in data from the dashed rectangle in a.

**Supplementary Figure 4:**
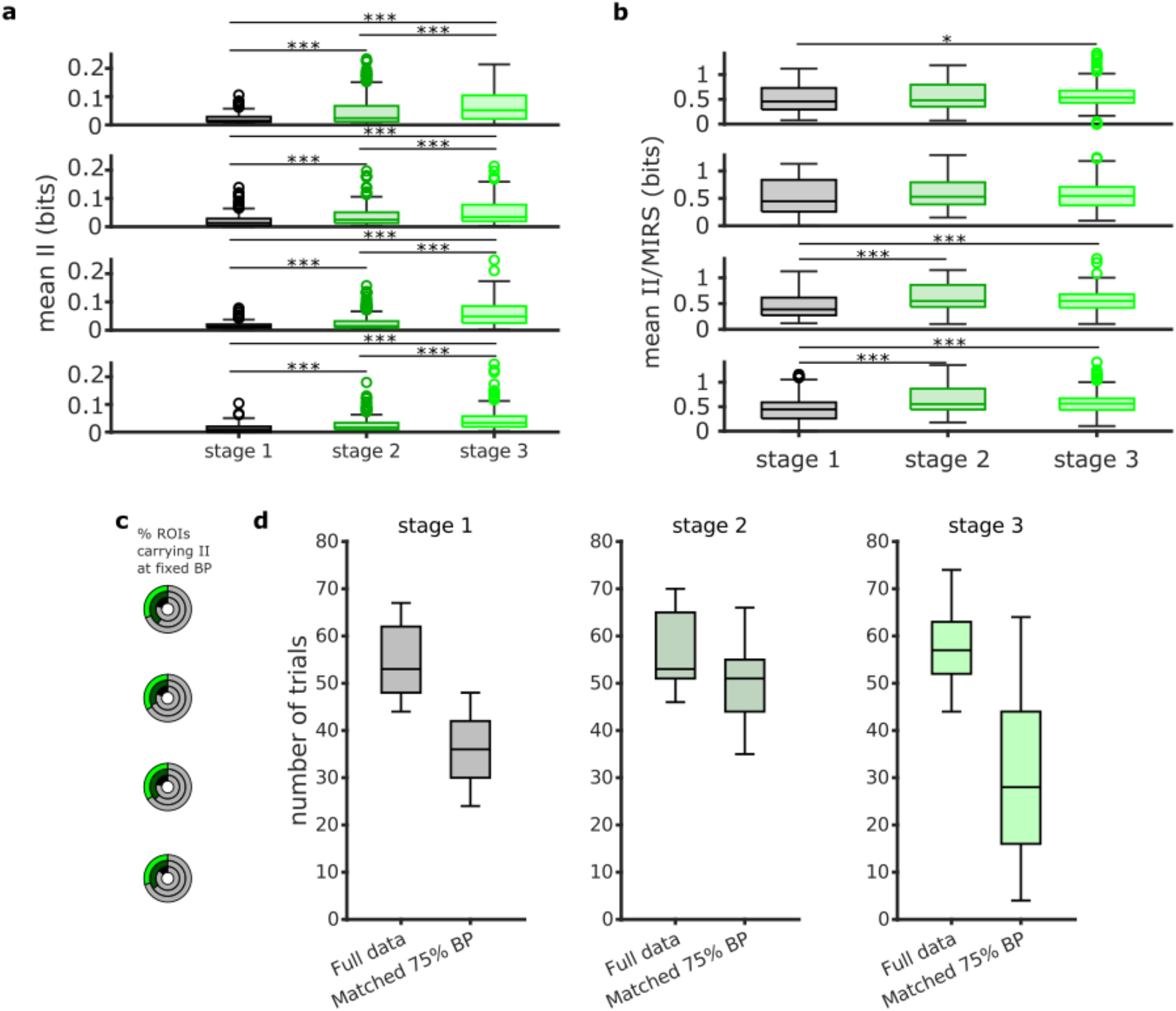
Increase in sensory readout efficacy in vS1 with sensory training does not depend on increased behavioral performance. **a**. Mean II across frames for all neurons with significant II for each learning stage and cortical depth when equalizing behavioral performance at 75% across all learning stages. Empty dots are outlier neurons. Horizontal lines indicate median value, and error bars indicate lower and upper quartiles. **b**. Same as in a, but for mean II/MIRS. **c**. Fraction of neurons carrying significant II at each cortical depth and at each learning stage when equalizing behavioral performance at 75% across all learning stages. Learning stage is indicated by the color inside each concentric circle. Full circles correspond to 100% of imaged neurons. The gray area in the circles indicates the fraction of neurons with non-significant II (p>=0.05). The remaining portion indicates neurons with significant II. **d**. Distributions of the number of available trials in all information-theoretical calculations (II, MIRS) in all ROIs, across all depths, when considering the original behavioral response of the animal (*Full data*) or when subsampling the trials to obtain a 75% behavioral performance (*Matched 75% BP*).

**Supplementary Figure 5:**
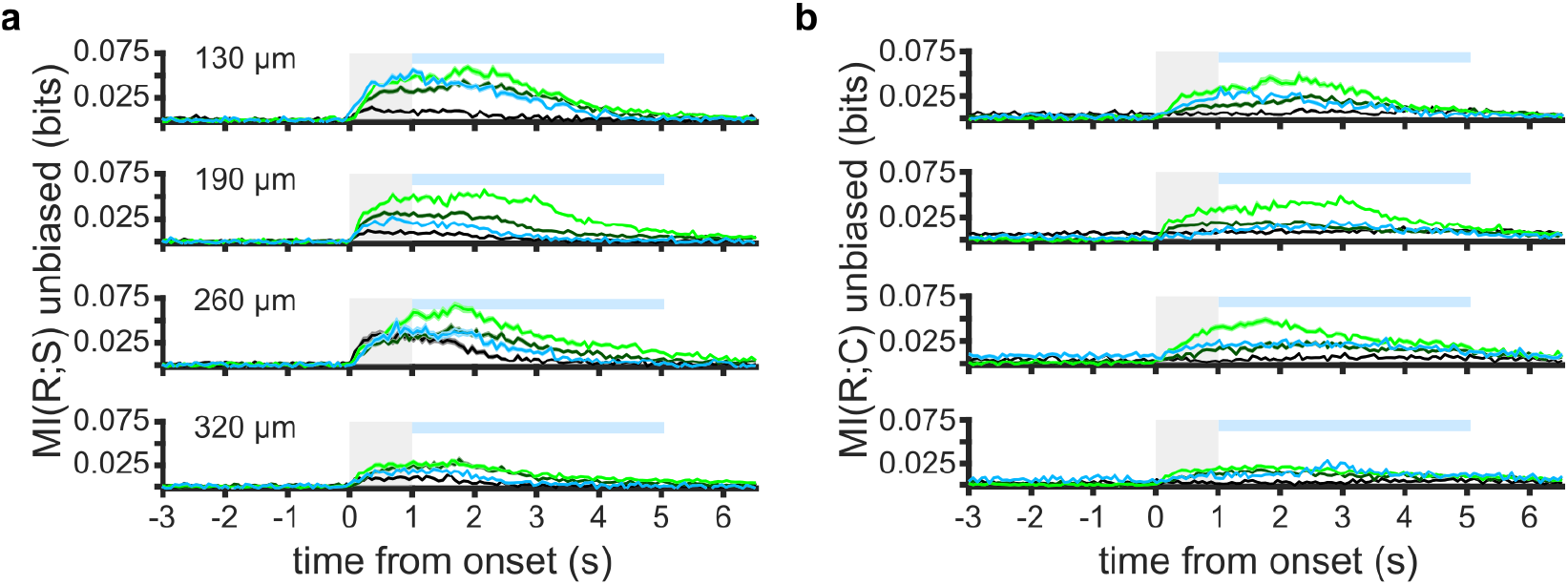
Comparison of MI-RS and MI-RC between stages 1-3 and switch session at all cortical depths. **a**. Mean frame-by-frame MI-RS across all neurons at each cortical depth and for each learning stage. Data are shown as in **Fig. 2c**, but only for the 3 mice used during the switch session. The gray shaded area indicates stimulus duration. The blue shaded area indicates the licking window. MI-RS was first averaged framewise across all neurons in the same FOV, and then averaged across all FOVs imaged at the same cortical depth and during the same learning stage. (stage 1: black, stage 2: dark green, stage 3: light green, switch session: blue). **b**. Same as in a., but for MI-RC.

**Supplementary Figure 6:**
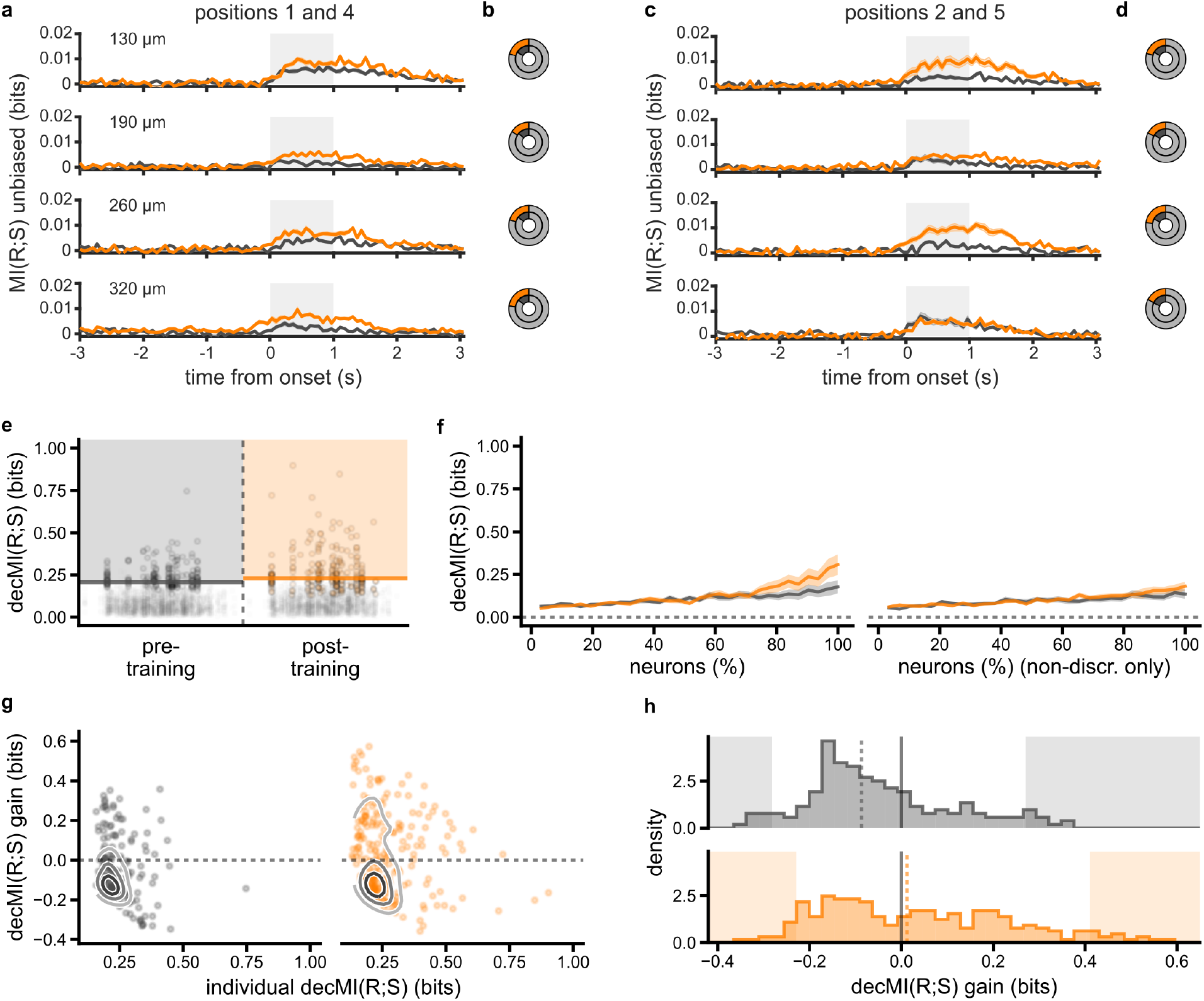
Sensory training improves general stimulus encoding in vS1. **a**. Mean MI-RS across all neurons imaged during the pre-training (gray) and post-training (orange) session, for each frame and for each cortical depth. MI-RS calculated on responses to stimulus positions 1 and 4 only. The gray shaded area indicates stimulus duration. **b**. Percentage of neurons carrying significant MI-RS (stimulus positions 1 and 4) for each imaging depth (p<0.05). Full circles indicate the total number of neurons. **c** and **d**, Same as a and b, but for stimulus positions 2 and 5. **e**. decMI-RS (balanced accuracy, stimulus decoding) plotted for individual ROIs pre-training and post-training (plotted with jitter in x). Discriminative neurons (decMI-RS p<0.05, compared to the chance decoding distribution) are plotted in dark gray (pre-training) or orange (post-training), and non-discriminative neurons are plotted in light gray (decMI-RS p>=0.05). Colored shading marks values above the 95th percentile. **f**. Mean across FOVs of population decMI-RS as neurons are added to the pool fed to the decoder, in order of lowest to highest individual decMI-RS. The full pool includes either all neurons (left) or only on-discriminative neurons (right) (pre-training: gray, post-training: orange). Stars indicate a significant difference pre to post training (t-test p<0.05). **g**. Discriminative neuron decMI-RS plotted against the gain in decMI-RS observed when decoders also received non-discriminative neuron responses as input. Contour lines qualitatively show data density levels. Pre-training and post-training are plotted left to right. **h**. Histograms of decMI-RS gain (y axis data in e), with pre-training and post-training plotted top to bottom. Zero gain is indicated by a black line, median gain is marked as the dashed line, and shaded areas show values below the 5th or above the 95th percentile of the distribution.

## Notes

### Competing Interest Statement

The authors have declared no competing interest.

## REFERENCES

1. Fanselow, E. E. & Nicolelis, M. A. L. Behavioral Modulation of Tactile Responses in the Rat Somatosensory System. J Neurosci 19, 7603–7616 (1999).

2. Pantoja, J. et al. Neuronal Activity in the Primary Somatosensory Thalamocortical Loop Is Modulated by Reward Contingency during Tactile Discrimination. J Neurosci 27, 10608–10620 (2007).

3. Chen, J. L. et al. Pathway-specific reorganization of projection neurons in somatosensory cortex during learning. Nat Neurosci 18, 1101–1108 (2015).

4. Yang, H., Kwon, S. E., Severson, K. S. & O’Connor, D. H. Origins of choice-related activity in mouse somatosensory cortex. Nat Neurosci 19, 127–134 (2016).

5. Bale, M. R., Bitzidou, M., Giusto, E., Kinghorn, P. & Maravall, M. Sequence Learning Induces Selectivity to Multiple Task Parameters in Mouse Somatosensory Cortex. Curr Biol 31, 473-485.e5 (2021).

6. Harrell, E. R., Renard, A. & Bathellier, B. Fast cortical dynamics encode tactile grating orientation during active touch. Sci Adv 7, eabf7096 (2021).

7. Rabinovich, R. J., Kato, D. D. & Bruno, R. M. Learning enhances encoding of time and temporal surprise in mouse primary sensory cortex. Nat Commun 13, 5504 (2022).

8. Buetfering, C. et al. Behaviorally relevant decision coding in primary somatosensory cortex neurons. Nat Neurosci 25, 1225–1236 (2022).

9. Chéreau, R. et al. Dynamic perceptual feature selectivity in primary somatosensory cortex upon reversal learning. Nat Commun 11, 3245 (2020).

10. Teich, A. F. & Qian, N. Learning and Adaptation in a Recurrent Model of V1 Orientation Selectivity. J Neurophysiol 89, 2086–2100 (2003).

11. Peron, S. P., Freeman, J., Iyer, V., Guo, C. & Svoboda, K. A Cellular Resolution Map of Barrel Cortex Activity during Tactile Behavior. Neuron 86, 783–799 (2015).

12. Petersen, R. S., Panzeri, S. & Diamond, M. E. Population Coding of Stimulus Location in Rat Somatosensory Cortex. Neuron 32, 503–514 (2001).

13. Panzeri, S., Moroni, M., Safaai, H. & Harvey, C. D. The structures and functions of correlations in neural population codes. Nat Rev Neurosci 23, 551–567 (2022).

14. O’Connor, D. H. et al. Vibrissa-Based Object Localization in Head-Fixed Mice. J Neurosci 30, 1947–1967 (2010).

15. O’Connor, D. H., Peron, S. P., Huber, D. & Svoboda, K. Neural Activity in Barrel Cortex Underlying Vibrissa-Based Object Localization in Mice. Neuron 67, 1048–1061 (2010).

16. Wekselblatt, J. B., Flister, E. D., Piscopo, D. M. & Niell, C. M. Large-scale imaging of cortical dynamics during sensory perception and behavior. J Neurophysiol 115, 2852–2866 (2016).

17. Chong, E. Z., Panniello, M., Barreiros, I., Kohl, M. M. & Booth, M. J. Quasi-simultaneous multiplane calcium imaging of neuronal circuits. Biomed Opt Express 10, 267 (2019).

18. Hromádka, T., DeWeese, M. R. & Zador, A. M. Sparse Representation of Sounds in the Unanesthetized Auditory Cortex. Plos Biol 6, e16 (2008).

19. Jadhav, S. P., Wolfe, J. & Feldman, D. E. Sparse temporal coding of elementary tactile features during active whisker sensation. Nat Neurosci 12, 792–800 (2009).

20. Barth, A. L. & Poulet, J. F. A. Experimental evidence for sparse firing in the neocortex. Trends Neurosci 35, 345–355 (2012).

21. Panzeri, S., Senatore, R., Montemurro, M. A. & Petersen, R. S. Correcting for the Sampling Bias Problem in Spike Train Information Measures. J Neurophysiol 98, 1064–1072 (2007).

22. Quiroga, R. Q. & Panzeri, S. Extracting information from neuronal populations: information theory and decoding approaches. Nat Rev Neurosci 10, 173–185 (2009).

23. Safaai, H., Heimendahl, M. von, Sorando, J. M., Diamond, M. E. & Maravall, M. Coordinated Population Activity Underlying Texture Discrimination in Rat Barrel Cortex. J Neurosci 33, 5843–5855 (2013).

24. Cooke, S. F. & Bear, M. F. Visual Experience Induces Long-Term Potentiation in the Primary Visual Cortex. J Neurosci 30, 16304–16313 (2010).

25. Frenkel, M. Y. et al. Instructive Effect of Visual Experience in Mouse Visual Cortex. Neuron 51, 339–349 (2006).

26. Polley, D. B., Steinberg, E. E. & Merzenich, M. M. Perceptual Learning Directs Auditory Cortical Map Reorganization through Top-Down Influences. J Neurosci 26, 4970–4982 (2006).

27. Law, C.-T. & Gold, J. I. Neural correlates of perceptual learning in a sensory-motor, but not a sensory, cortical area. Nat Neurosci 11, 505–513 (2008).

28. Pica, G. et al. Quantifying how much sensory information in a neural code is relevant for behavior. in Advances in Neural Information Processing Systems vol. 30 (Curran Associates, Inc., 2017).

29. Panzeri, S., Piasini, E. & Fellin, T. Cracking the Neural Code for Sensory Perception by Combining Statistics, Intervention, and Behavior. Neuron 93, 491 507 (2017).

30. Zuo, Y. et al. Complementary Contributions of Spike Timing and Spike Rate to Perceptual Decisions in Rat S1 and S2 Cortex. Curr Biol 25, 357–363 (2015).

31. Valente, M. et al. Correlations enhance the behavioral readout of neural population activity in association cortex. Nat Neurosci 24, 975–986 (2021).

32. Saleem, A. B., Diamanti, E. M., Fournier, J., Harris, K. D. & Carandini, M. Coherent encoding of subjective spatial position in visual cortex and hippocampus. Nature 562, 124–127 (2018).

33. Chen, J. L., Carta, S., Soldado-Magraner, J., Schneider, B. L. & Helmchen, F. Behaviour-dependent recruitment of long-range projection neurons in somatosensory cortex. Nature 499, 336–340 (2013).

34. Poort, J. et al. Learning Enhances Sensory and Multiple Non-sensory Representations in Primary Visual Cortex. Neuron 86, 1478–1490 (2015).

35. Francis, N. A. et al. Small Networks Encode Decision-Making in Primary Auditory Cortex. Neuron 97, 885-897.e6 (2018).

36. Stringer, C. et al. Spontaneous behaviors drive multidimensional, brainwide activity. Science 364, eaav7893 (2019).

37. Francis, N. A. et al. Sequential transmission of task-relevant information in cortical neuronal networks. Cell Reports 39, 110878 (2022).

38. Tseng, S.-Y., Chettih, S. N., Arlt, C., Barroso-Luque, R. & Harvey, C. D. Shared and specialized coding across posterior cortical areas for dynamic navigation decisions. Neuron 110, 2484-2502.e16 (2022).

39. Keller, G. B., Bonhoeffer, T. & Hubener, M. Sensorimotor Mismatch Signals in Primary Visual Cortex of the Behaving Mouse. Neuron 74, 809--815 (2012).

40. McGuire, L. M. et al. Short Time-Scale Sensory Coding in S1 during Discrimination of Whisker Vibrotactile Sequences. Plos Biol 14, e1002549 (2016).

41. Pala, A. & Stanley, G. B. Ipsilateral Stimulus Encoding in Primary and Secondary Somatosensory Cortex of Awake Mice. J Neurosci 42, 2701–2715 (2022).

42. Schoups, A., Vogels, R., Qian, N. & Orban, G. Practising orientation identification improves orientation coding in V1 neurons. Nature 412, 549–553 (2001).

43. Kato, H. K., Gillet, S. N. & Isaacson, J. S. Flexible Sensory Representations in Auditory Cortex Driven by Behavioral Relevance. Neuron 88, 1027–1039 (2015).

44. Banerjee, A. et al. Value-guided remapping of sensory cortex by lateral orbitofrontal cortex. Nature 585, 245–250 (2020).

45. Feldman, D. E. Timing-Based LTP and LTD at Vertical Inputs to Layer II/III Pyramidal Cells in Rat Barrel Cortex. Neuron 27, 45–56 (2000).

46. Bender, V. A., Bender, K. J., Brasier, D. J. & Feldman, D. E. Two Coincidence Detectors for Spike Timing-Dependent Plasticity in Somatosensory Cortex. J Neurosci 26, 4166–4177 (2006).

47. Voelcker, B., Pancholi, R. & Peron, S. Transformation of primary sensory cortical representations from layer 4 to layer 2. Nat Commun 13, 5484 (2022).

48. Margolis, D. J. et al. Reorganization of cortical population activity imaged throughout long-term sensory deprivation. Nat Neurosci 15, 1539–1546 (2012).

49. Carrillo-Reid, L., Han, S., Yang, W., Akrouh, A. & Yuste, R. Controlling Visually Guided Behavior by Holographic Recalling of Cortical Ensembles. Cell 178, 447-457.e5 (2019).

50. Dalgleish, H. W. et al. How many neurons are sufficient for perception of cortical activity? Elife 9, e58889 (2020).

51. Gill, J. V. et al. Precise Holographic Manipulation of Olfactory Circuits Reveals Coding Features Determining Perceptual Detection. Neuron 108, 382-393.e5 (2020).

52. Heffner, H. E. & Heffner, R. S. Hearing ranges of laboratory animals. J Am Assoc Laboratory Animal Sci Jaalas 46, 20–2 (2007).

53. Linden, J. F., Liu, R. C., Sahani, M., Schreiner, C. E. & Merzenich, M. M. Spectrotemporal Structure of Receptive Fields in Areas AI and AAF of Mouse Auditory Cortex. J Neurophysiol 90, 2660–2675 (2003).

54. Akam, T. et al. Open-source, Python-based, hardware and software for controlling behavioural neuroscience experiments. Elife 11, e67846 (2022).

55. Guo, Z. V. et al. Flow of Cortical Activity Underlying a Tactile Decision in Mice. Neuron 81, 179–194 (2014).

56. Juavinett, A. L., Nauhaus, I., Garrett, M. E., Zhuang, J. & Callaway, E. M. Automated identification of mouse visual areas with intrinsic signal imaging. Nat Protoc 12, 32–43 (2017).

57. Pachitariu, M. et al. Suite2p: beyond 10,000 neurons with standard two-photon microscopy. Biorxiv 061507 (2017) doi:10.1101/061507.

58. Chen, T.-W. et al. Ultra-sensitive fluorescent proteins for imaging neuronal activity. Nature 499, 295–300 (2013).

59. Panzeri, S. & Treves, A. Analytical estimates of limited sampling biases in different information measures. Netw Comput Neural Syst 7, 87–107 (2018).

60. Optican, L. M. & Richmond, B. J. Temporal encoding of two-dimensional patterns by single units in primate inferior temporal cortex. III. Information theoretic analysis. J Neurophysiol 57, 162–178 (1987).

61. Ince, R. A. A., Mazzoni, A., Bartels, A., Logothetis, N. K. & Panzeri, S. A novel test to determine the significance of neural selectivity to single and multiple potentially correlated stimulus features. J Neurosci Meth 210, 49–65 (2012).

62. Bertschinger, N., Rauh, J., Olbrich, E., Jost, J. & Ay, N. Quantifying Unique Information. Entropy 16, 2161–2183 (2014).

63. Strong, S. P., Koberle, R., Steveninck, R. R. de R. van & Bialek, W. Entropy and Information in Neural Spike Trains. Phys Rev Lett 80, 197–200 (1998).

64. Magri, C., Whittingstall, K., Singh, V., Logothetis, N. K. & Panzeri, S. A toolbox for the fast information analysis of multiple-site LFP, EEG and spike train recordings. Bmc Neurosci 10, 81 (2009).

65. Makkeh, A., Theis, D. O. & Vicente, R. BROJA-2PID: A Robust Estimator for Bivariate Partial Information Decomposition. Entropy 20, 271 (2018).

66. Harris, C. R. et al. Array programming with NumPy. Nature 585, 357–362 (2020).

67. Virtanen, P. et al. SciPy 1.0: fundamental algorithms for scientific computing in Python. Nat Methods 17, 261–272 (2020).

68. McKinney, W. Data Structures for Statistical Computing in Python. in Proceedings of the 9th Python in Science Conference SCIPY 2010 vol. 445 51–56 (2010).

69. Hunter, J. D. Matplotlib: A 2D Graphics Environment. Comput Sci Eng 9, 90–95 (2007).

70. Pedregosa, F. et al. Scikit-learn: Machine Learning in Python. Journal of Machine Learning Research 12, (2011).

